# Application of a high molecular weight topical hyaluronan formulation blocks keratinocyte tumor initiation downstream of UVB-induced oncogenic mutations

**DOI:** 10.64898/2026.11.24.768908

**Authors:** C. Tolg, M. Liu, K. Cousteils, H. Pavanel, T.A. Miller, C.S. Taylor, D. Chen, K.A. Hill, E.A. Turley

**Affiliations:** Dept. Biochemistry, Schulich School of Medicine and Dentistry, Western University, London ON CA; Dept. Biology, Faculty of Science, Western University, London ON CA

## Abstract

The polysaccharide hyaluronan in its high molecular weight form (HMW-HA) has been implicated as providing tumor resistance. Since HMW-HA is depleted with age and UVB exposure, we directly assessed the effect of replacing HMW-HA in UVB-exposed skin of *Hr*^-/-^:*Ptch1*^LacZ/+^ mice, a model of susceptibility to keratinocyte tumors. We developed a fragmentation-resistant, topical HMW-HA formulation (HMW-HA-phosphatidylethanolamine, HAPE) that penetrates skin to form coats around keratinocytes, which are not depleted by UVB. HAPE prevents detectable UVB-induced tumors and promotes epidermal homeostasis even though whole exome sequencing shows that the UVB-induced single base pair mutation burden is similar to control. Mutagenesis of most keratinocyte driver oncogenes is not affected, but mutation of *Trp53*, which is required for sustained keratinocyte tumorigenesis, is rarely detected in HAPE-treated skin. Immunofluorescence and spatial transcriptomics (10X genomics) show that HAPE blunts expression of oncogenic driver pathways (e.g. sonic hedgehog) throughout the epidermis but selectively reduces TRP53 protein expression and signaling in hair follicles. Spatial transcriptomics identifies a hair follicle KRT15+:TRP53+ stem cell subset that is lost with HAPE treatment. Since KRT15+ hair follicle progenitors are cancer-initiating, we assessed the effect of HMW-HA on the tumorigenesis of human keratinocytes bearing gain-of-function TRP53 mutations. HMW-HA promotes apoptosis of KRT15+ subsets, prevents disorganized and invasive growth of xenografts and supports polarized keratinocyte architecture. These results identify a novel microenvironmental tumor suppression mechanism that selectively targets mutant cancer-initiating cells and maintains skin homeostasis and polarity.

## INTRODUCTION

Previous reports identified HMW-HA as a cancer-resistance factor in naked mole rats^[1]^, which are long-lived (30 yrs.) rodents that rarely develop cancers, particularly in skin^[2,3^]. The transformation-resistance of naked mole rat skin fibroblasts by oncogenes *in vitro*^[1]^ depends on HMW-HA. Cutaneous squamous cell carcinoma, (cSCC) formed in response to chemical carcinogens are reduced in transgenic C57BL/6 mice expressing naked mole rat HA synthase 2^[4]^, resulting in accumulation of HMW-HA. To date, the tumor-resistance of naked mole rat HA is attributed in part to the anti-inflammatory properties of its very high molecular weight form, and reduced hyaluronidase expression in naked mole rat skin^[5]^. HMW-HA also promotes apoptosis in cancer cell lines, which is an event linked to regulating the expression of mutant TP53^[6]^. These reports and evidence that HMW-HA is reduced in skin with age and UVB exposure^[7–10]^ coincident with increased tumor incidence, prompted us to both assess if this polysaccharide alters UVB-induced keratinocyte tumor initiation, and to probe HMW-HA anti-tumor functions.

We chose to assess the effect of HMW-HA on keratinocyte tumors because these are the most common cancers worldwide and are a large health care burden^[11]^. They are generally not lethal but are clinically important because of their increasing incidence, targeted treatment resistance and frequent recurrence^[12^]. Current treatments, which include targeted and non-specific approaches have not been particularly effective in reducing recurrence^[13,14^]. The major carcinogenic cause of these tumors is UV exposure (https://www.cancer.gov/types/skin). Tissues that are chronically exposed to mutagens/carcinogens such as UVB develop large areas of somatic mutations that increase the risk of multiple, and recurrent tumors^15,16^]. Ultradeep, targeted DNA sequencing has shown that histologically normal, healthy skin of aged donors exposed to sun unexpectedly contains ‘driver’ somatic mutations in cancer-causing genes^15,17, 18^] with 1 in every 4 cells predicted to carry an oncogenic mutation^17^]. These results have been used as evidence that somatic driver mutations are a first step toward cancer initiation, but additional downstream factors are required to achieve stable transformation. For example, the composition of the extracellular matrix, including HA, can profoundly regulate oncogene-mediated tumor initiation^19^].

We used a UVB-induced cutaneous squamous and basal cell carcinomas in a *Hr*^-/-^:*Ptch1*^+/-^ mouse model of susceptibility to probe HMW-HA functions that play a role in restraining keratinocyte cancers induced by UVB^20,21^]. This model is ideal for this analysis since it mimics human keratinocyte tumors induced by exposure to UV by generating both squamous cell and basal cell carcinomas from a large area of field mutations. Importantly, the cancer initiating cells have been identified in this model as KRT15^+^ or KRT14^+^ hair follicle subsets^20–22^] allowing an assessment of their role in any anti-cancer properties of HMW-HA.

Our experimental approach was to replace HMW-HA lost with chronic UVB exposure using a topical HMW-HA phosphatidylethanolamine co-polymer formulation (HAPE) that, unlike unmodified HMW-HA, penetrates the epidermis to form coats around keratinocytes^23^]. We used multi-omics to probe the effects of HAPE and show that HAPE prevents the initiation of cutaneous basal and squamous cell carcinomas by at least two mechanisms. It blocks activation of oncogenic driver pathways such as Sonic Hedgehog (SHH) signaling downstream of UVB-induced mutations, and targets subpopulations of KRT15 cancer-initiating hair follicle cells bearing mutant TRP53 to restrict their survival.

## METHODS

### Animal Housing

Mice were individually caged in a temperature-controlled environment with a twelve-hour light/dark cycle and fed PicoLab mouse diet (LabDiet, St. Louis, MO, United States) ad libitum. All experiments were approved by and compliant with the standard operating procedures of the Animal Use Subcommittee at the University of Western Ontario, Canada (2009-060). Female *Hr*^-/-^/SKH-1 mice (Strain 477, Charles River Laboratories, Wilmington, MA, USA) were crossed with *Ptch*^+/LacZ^/C57BL/6 male, haired mice (Stock number 003091, Jackson Laboratory, Bar Harbor, ME, USA). F1 offspring between 14 to 21 days of age were genotyped for *Ptch^+/-^*heterozygosity. F1 *Ptch^+/-^/Hr^+/-^* male mice were crossed with F1 *Ptch^+/-^/Hr^+/-^* female mice, and F2 *Ptch^+/-^/Hr^-/-^* mice that were both hairless (*Hr^-/-^*) and *Ptch^+/-^* were utilized for the study.

### Tail DNA Isolation

When mice were between 14 and 21 days of age, tissue samples smaller than 0.5 cm were taken from the tails of mice and stored at -20°C until genotyping was performed. Tissue was lysed overnight at 55°C using 350 μL lysis buffer (1 M Tris, 0.5 M EDTA, 10% SDS, 5 M NaCl, pH 8) and 300 μg proteinase K (Roche, Mannheim, Germany). The following day, samples were spun at 208 x g for 10 minutes at room temperature. The supernatant was transferred to new tubes containing 500 μL isopropanol. After mixing, samples were spun at 208 x g for 10 minutes at room temperature. The supernatant was discarded, and the pellet was washed with 500 μL of ethanol (−20°C). Tubes were spun at 208 x g for 10 minutes at room temperature, washed with ethanol (−20°C) again, and then spun at 208 x g for 5 minutes at room temperature. Supernatant was discarded, and DNA pellets were left to air dry at room temperature overnight. DNA was resuspend in 100 μL of ddH_2_O and incubated at 55°C for 15 minutes to dissolve the pellet. DNA concentration was determined using NanoDrop (ThermoScientific, Waltham, MA, United States). The ratio of the absorption at 260 nm over the absorption at 280 nm varied between 1.85 to 1.97. DNA concentration was adjusted to 50 ng/μL using ddH_2_O.

### Genotyping

Genotyping to confirm *Ptch1* heterozygosity was performed using the protocol provided on the Jackson Website (“Genotyping Protocol for Stock Number: 003081,” 2010) and outlined below. Primers (Invitrogen, Waltham, MA, United States) were dissolved in ddH_2_O to a concentration of 100 nmoles/mL. For the PCR reaction, each tube contained 12.5 μL AmpliTaq Gold (ThermoFisher, Waltham, MA, United States), 1 μL of GC Enhancer (ThermoFisher, Waltham, MA, United States), 0.25 μL of each 100 nmole/mL primer solution, and 8.5 μL of ddH_2_O. To each tube 2 μL of tail DNA was added.

**Table 1.**
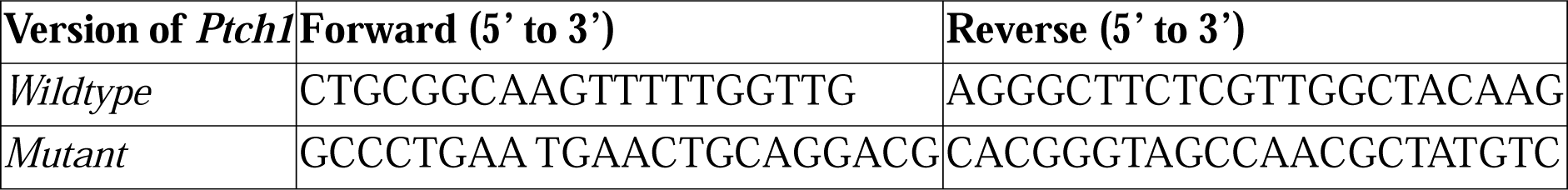
Primers for *Ptch1*.

The PCR reaction was done as follows: 94°C for 1.5 minutes to start; cycle through 94°C for 30 seconds, for 30 seconds, 69°C minute, 72°C for 45 seconds, and repeat 35 times; finally, hold at 10°C. Amplified samples were separated by gel electrophoresis on a 1.5% agarose gel in 1X TBE (90 mM Tris-borate, 2 mM EDTA, pH 8.3) buffer containing 50 μL of a 1 mg/mL aqueous solution of ethidium bromide. 10x DNA gel-loading dye (3.9 mL glycerol, 500 μL 10% (w/v) SDS, 200 μL 0.5 M EDTA and 0.025 g bromophenol blue in 10 mL) was added to PCR samples and DNA ladder (ThermoFisher, Waltham, MA, United States) at a 10:1 ratio prior to loading. Gel electrophoresis occurred at 110 V for 1 hour. The gel was imaged using the Molecular Imager® Gel Dock TM XR+ with ImageLab TM Software (Universal Hood II, Bio-Rad Laboratories, Hercules, California, United States). Under these conditions, the wild-type *Ptch1* allele forms a band of 200 base pairs in length, and the mutant allele forms a band of 479 base pairs in length.

### Preparation of HAPE

HAPE cream (PCT CA270352, Turley, 2011)^23^] was prepared as previously described^23^]. Briefly, 1.35 mL of 1% (v/w) Sodium Hyaluronate Solution (500 kDa, Medical Grade, Lifecore Biomedical, Chaska, MN, United States) and 252 L of isopropanol was mixed thoroughly with 1.35 mL of unrefined liquid soy lecithin (Soy Lecithin GT non-GM IP, Imperial-Oel-Import, Germany) at room temperature. Dry (3.78 mg) 1-ethyl-3-(3-dimethylaminopropyl) carbodiimide (EDC) (Sigma, St. Louis, MO, United States) was added and mixed thoroughly. After mixing for 10 to 15 minutes, 3 mL of vehicle cream was added and mixed for an additional 5 minutes. A water-based cream (Mango Face cream, Aquatech, Mississauga, Ontario, Canada) was used throughout the study as the vehicle for mixing with HAPE or unaltered as the vehicle cream control. Both the HAPE and vehicle creams were stored protected from light at 4 .

### HAPE Treatment of *Hr*^-/-^:*Ptch1*^+/-^ mice

*Hr^-/-^/ Ptch^+/-^* mice were separated into four experimental groups: No UVB control (No UVB, n= 5), UVB-only control (UVB, n=7), UVB with vehicle cream control (UVB + vehicle, n=10), and UVB with HAPE cream (UVB+HAPE, n=10). An UV-irradiation unit (Daavlin Co., Bryan, OH, USA) equipped with an electronic UV-dosage controller^186^ was used to administer UV radiation. Mice between 6 to 7 weeks of age were exposed to UV irradiation consisting of UVA of 315-400 nm, accounting for 20% of total energy; and UVB of 290-315 nm, accounting for 80% of total energy. Mice received 180 mJ/cm^2^ of UVB irradiation twice per week (Thursdays and Fridays) for 30 to 45 seconds, while vehicle or HAPE cream was applied prior to UVB-irradiation, once daily for five days each week (Monday to Friday). Mice were irradiated for 4 weeks and 26 weeks to study the acute and chronic effects of HAPE on UVB-induced BCC, respectively. Lesions and skin abnormalities were measured using digital calipers (500-170-30, Aurora, IL, USA), excised using 8-mm punch biopsy from the dorsal back, and processed to be paraffin-embedded for histological analyses. Lesion-adjacent phenotypically normal skin was also excised using an 8-mm punch biopsy, frozen using 70% ethanol and dry ice and stored at -80 . Histopathology analysis of skin abnormalities was conducted at the Centre for Phenogenomics (Toronto, ON, CA) using paraffin-embedded slides stained with H&E. Neoplasia was identified by histological features characteristic of BCC (large nuclei, scant cytoplasm, mitotic features), cell morphology, and clonal expansion.

### Immunohistochemistry

Immunohistochemistry of paraffin-embedded tissue was conducted as previously described^193^. Briefly, sections were de-paraffinized through two passages of xylene for 15 minutes each wash and rehydrated through a descending ethanol series (100%, 95%, 70%) for 10 minutes each followed by ddH_2_O for 5 minutes. Antigen retrieval was performed using 10mM sodium citrate buffer (pH 6.0) in a standard microwave. Specimen were washed in two changes of the appropriate buffer for 5 minutes each, and endogenous peroxidase activity was inhibited by incubating slides in 3% H_2_O_2_ for 20 minutes. Non-specific antibody binding was blocked by incubating slides in 3% bovine serum albumin (BSA) at room temperature for 1 hour. Diluted primary antibody was added to each slide and incubated at 4 overnight in humidifying chambers, with non-immune anti-mouse immunoglobulin (IgG) incubated on the negative control slide. Following two washes in the appropriate buffer solution, the slides were incubated with 1:100 biotinylated goat anti-rabbit secondary antibody (catalog number: E0432, Agilent Technologies, CA, USA) diluted in phosphate buffer saline (PBS: 137 mM NaCl, 12 mM phosphate, 2.7 mM KCl, pH 7.4) or Tris-buffer saline (TBS: 50 mM Tris-Cl, pH 7.5, 150 mM NaCl) at room temperature for 2 hours in humidifying chambers. Streptavidin-conjugated horseradish-peroxidase (catalog number: ab7403, AbCam, MA, USA) was diluted in 1xPBS or TBS to 1:1000-fold and added onto each slide for 30 minutes at room temperature, followed by incubation with diaminobenzidine substrate (DAB, catalog number: K3467, Agilent Technologies, CA, USA). Counterstaining was carried out using hematoxylin diluted 1:10-fold in ddH_2_O (10143-606, VWR, PA, USA). Sections were dehydrated by passaging through the ethanol series (70%, 95%, 100%) and two changes of xylene. All slides were mounted using Cytoseal-60 (catalog number: 8310-16, ThermoFisher Scientific, MA, USA) and dried at room temperature.

### Image analysis of histochemistry

All sections were scanned using Aperio ScanScope (Leica Biosystems, Wetzlar, Germany) and analyzed using ImageJ (https://downloads.imagej.net/fiji/) or QuPath^24^]. All quantifications were conducted on the epidermis of animal sections, and keratinocytes within the field of view were quantified according to their epidermal compartmentalization (supra-basal layer, stratum basale, and hair follicles). For intensity quantification, the DAB channel in each image was separated from the hematoxylin channel using the “Colour deconvolution” algorithm and measured using the “Pixel measurement” algorithms in ImageJ. DAB-stained brown pixels in slides from all experimental groups were compared to the negative control, which was used as the threshold. Semi-automatic positive cell counts of all slides were conducted using the “Positive cell detection” algorithm from QuPath (v0.1.9) as previously described^24^] and all algorithm-generated data were verified by manual counting. Immunofluorescence and image analysis. Immunofluorescence of paraffin-embedded tissue sections was conducted identically to the protocol for immunohistochemistry described above, with the quenching of endogenous peroxidase omitted. After secondary antibody incubation using either Alexa 488-conjugated mouse anti-rabbit IgG (catalog number: A32731, Invitrogen, MA, USA) at 1:200-fold dilution or Alexa 555-conjugated mouse anti-mouse IgG (catalog number: A32727, Invitrogen, MA, USA) at a 1:200-fold dilution, slides were washed in the appropriate buffer solution. Sections were then either mounted using DAPI-containing ProLong Gold anti-fade mounting reagent (catalog number: P36931, ThermoFisher Scientific, MA, USA) or stained with DAPI and mounted with ProLong Gold anti-fade mounting reagent (catalog number: P36930, ThermoFisher Scientific, MA, USA). All tissue sections were imaged using IX81 Olympus confocal microscope equipped with FV10-ASW 4.2 software at 20x or 40x magnifications. Images were saved in TIFF format and quantified using ImageJ or QuPath as described above. Keratinocytes within the field of view were quantified according to their epidermal compartmentalization (suprabasal layer, stratum basale, and hair follicles). To quantify the intensity of antibody immunoreactivity, DAPI, FITC and TexasRed channels were first separated in ImageJ using the “Split channel” algorithm, and threshold against the negative control. Epidermal compartments in each field of view were selected manually, and the “Pixel measurement” algorithm was used to detect intensity. Similarly, individual colour channels were separated in QuPath and the “Positive cell detection” algorithm was used to detect keratinocytes positive for antibody staining and all algorithm-generated counts were verified manually.

### HaCaT cell culture

HaCaT keratinocytes were the kind gift of Dr. L. Laird (Dept. Anatomy and Cell Biology, Western University, London ON CA). Cells were cultured in DMEM high glucose medium containing 10% FBS and antibiotics/antimycotics at 37°C, H_2_O saturated atmosphere. When adherent cultures were 80% confluent, cells were passaged using Trypsin/EDTA. For immunostaining, cells were seeded at subconfluence on cover slips using culture conditions described above.

### Particle exclusion assay

24 hours after plating HaCaT keratinocytes, 750 μL of a suspension of fixed sheep erythrocytes (≈10^8^/mL) was added to the cells and allowed to settle for 15 minutes. Hyaluronan coat formation was documented using phase contrast microscopy. Cells were treated with hyaluronan-specific *Streptomyces* hyaluronidase at 1 U/mL of medium for 1 hour before visualization with the particle assay, to demonstrate the structural dependence of clear coat area as dependent upon hyaluronan.

### RNA extraction and *in vitro* reverse transcription

RNA extraction from tissue was performed with trizol/chloroform extraction protocol. All equipment was cleaned with 70% ethanol and RNase-away solution (catalog number: 10328-011, Invitrogen, MA, USA). Complementary DNA (cDNA) was synthesized using SuperScript IV VILO Master Mix kit (catalog number: 11756050, Invitrogen, MA, USA). 4 μL of SuperScript IV VILO master mix solution was incubated with 1 μg of RNA template and nuclease-free water for a final reaction volume of 20 μL. Samples were gently mixed and incubated at 25 for 10 minutes to allow primer annealing, followed by incubation at 50 for 1 hour to allow elongation. Samples were then heated to 85 for 5 minutes to inactivate reverse transcriptase, briefly centrifuged and stored at -20 until further assays were performed.

### PCR reactions and analysis

RT-PCR was performed to analyze *CD44* expression in transgenic mice. Each PCR reaction consisted of 20 ng of cDNA template, 0.5 μL of 10 μM forward and reverse primers (Table 2), 0.5 μL of GC enhancer, 10 μL of AmpliTaq 360 master mix, and nuclease-free ddH_2_O to reach a total volume of 25 μL. PCR reactions were carried out at 95 for 10 minutes, 94 for 30 seconds, 55 for 30 seconds, 72 for 30 seconds for 40 cycles, and a final elongation step at 72 for 7 minutes using DNA Engine Tetrad 2 Thermocycler (Bio Rad, CA, USA). All target gene expression was compared with the housekeeping control gene GAPDH on a 1% agarose gel containing ethidium bromide, and imaged using ChemiDoc^TM^ XRS+ System (Bio Rad, CA, USA).

**Table 2:**
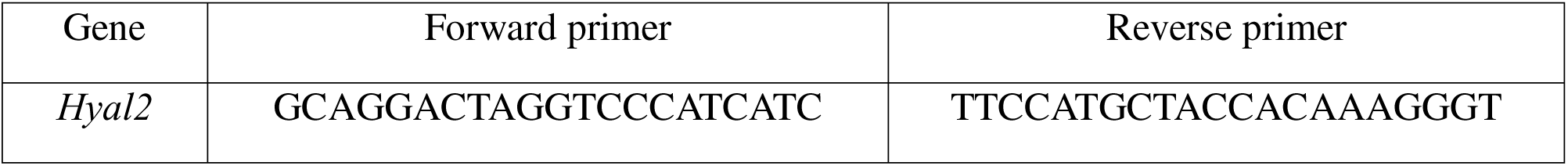
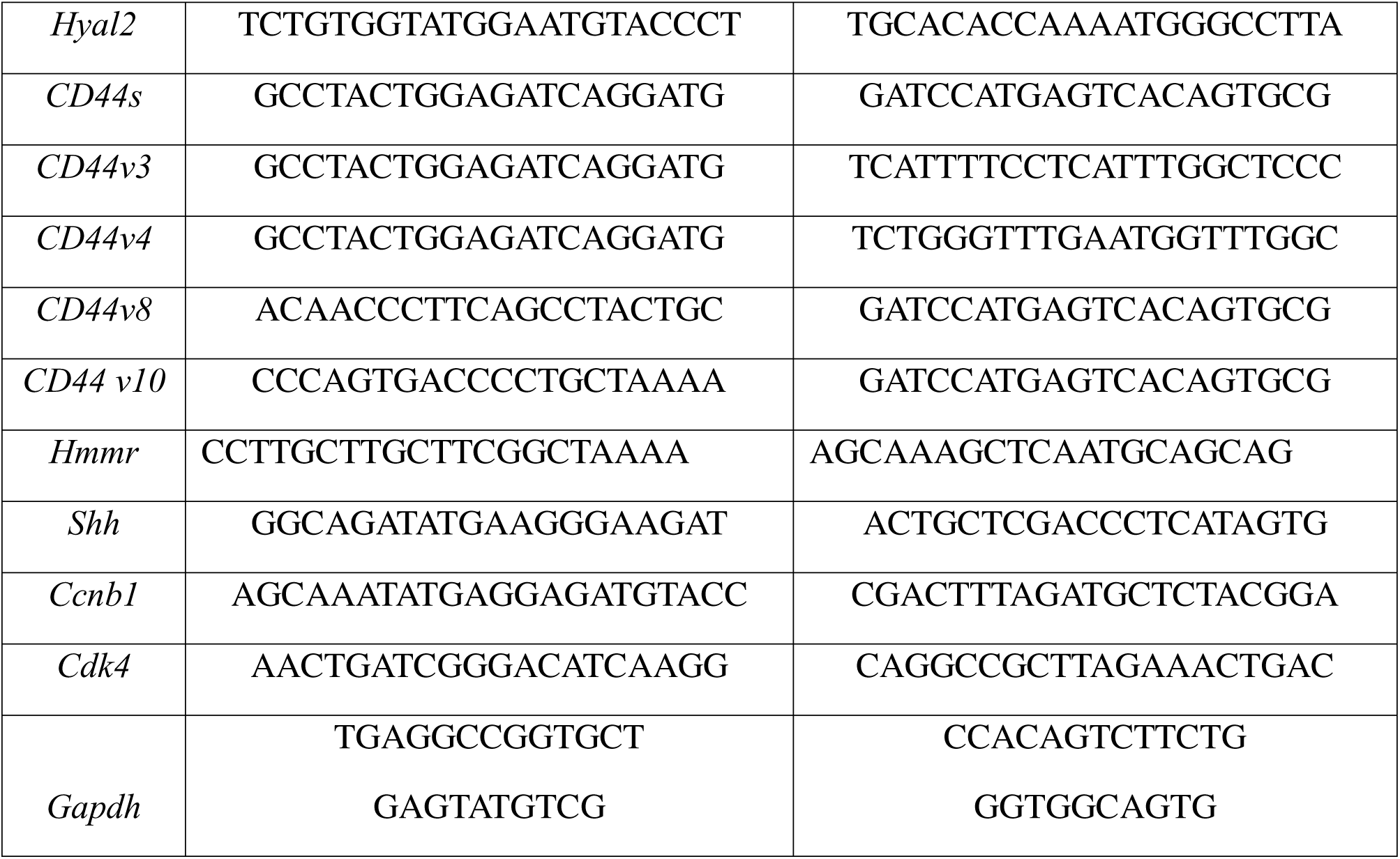
Primers used in RT-qPCR and RT-PCR.

### Real-time PCR reactions and analysis

RT-qPCR was performed to analyze the mRNA expression of *Hyal2, Hyal3, Hmmr, Shh, Cccnb1, Cdk4,* and *Gapdh* in transgenic mice. Each PCR reaction consisted of 20 ng of cDNA template, 0.5 uL of 10 uM forward and reverse primers (Table 2), 10 μL of SYBR Green PCR Master Mix (catalog number: 4309155, Thermofisher Scientific, MA, USA), and nuclease-free ddH_2_O to reach a total volume of 20 μL. PCR reactions were carried out on a 96-well plate at 95 for 3 minutes, 60 for 10 seconds, and 72 for 30 seconds for 40 cycles using Stratagene Mx3000P system (Agilent Technologies, Santa Clara, CA). Ct values were exported from the thermocycler and analyzed using Microsoft excel according to the 2^-△CT^ method.

### DNA extraction from tissue

Approximately 1 g of frozen tissue sample was incubated in lysis buffer (1M Tris, 0.5M EDTA, 10% SDS, 5M NaCl, pH 8.0) and 20 mg/mL proteinase K (Roche, Mannheim, Germany) overnight at 55 with constant agitation. Samples were centrifuged at 200xg for 10 minutes at room temperature, and the supernatants in each sample were transferred into new tubes. 100 μL of protein precipitation solution (catalog number: A975A, Promega Corporation, WI, USA) was added to samples and centrifuged at 14,000xg for 10 minutes. Supernatants were briefly chilled on ice followed by centrifugation at 14,000xg for 5 minutes, with pellets discarded. Immediately after, ice-cold isopropanol (500 μL) was added to supernatants, mixed by inversion five times and incubated at -20 for 30 minutes. All samples were centrifuged at 14,000xg for 15 minutes, and pellets were washed with ice-cold 70% ethanol. Centrifugation and wash step were repeated. The resulting pellets were air-dried at room temperature and suspended in 20 μL of ddH_2_O. Sample concentration was measured using NanoDrop One/One^C^ (ThermoFisher Scientific, MA, USA) and stored in -20 until further use. Only samples with an A_260/280_ ratio between 1.8 to 1.9 were used in future experiments.

### 8-oxo-2’-deoxyguanosine ELISA

Nuclear ROS-adducts 8-oxo-2’-deoxyguanosine (8-oxo-dG) were detected using a commercially available ELISA kit (catalog number: 4380-096-K, Trevigen, MD, USA). DNA was extracted from frozen tissue as described above. Samples were diluted in nuclease-free ddH_2_O to 200 μg/mL, and 100X cations (catalog number: 4380-096-05) was added for a final 1X concentration. 0.8 μL of DNase I (catalog number: 4380-096-06, 5 U/uL) and alkaline phosphatase (catalog number: 4386-096-07) were added to each sample and incubated at 37 for 1 hour separately. Digested samples were aliquoted and stored at -20 until assayed. 8-oxo-dG standards (catalog number: 4380-096-01, 20 μM) were serially diluted with assay diluent (catalog number: 4380-096-02) to construct a standard curve (200 nM, 100 nM, 50 nM, 25 nM, 12.5 nM, 6.25 nM, and 3.13 nM). 25 μL of standards, blank and DNA samples were pipetted in triplicates into a 96-well plate pre-coated with 8-oxo-dG, with 25 μL of anti-8-OHdG monoclonal antibody (catalog number: 4380-096-03, diluted 1:250-fold in assay diluent) added to each well. The plate was covered with a film sealer and incubated for 1 hour at 25 . All wells were washed with 4 changes of 1xPBST (137 mM NaCl, 12 mM phosphate, 2.7 mM KCl, pH 7.4, 0.1% Tween-20) for 30 seconds, with excess liquid aspirated. Goat-anti-mouse IgG-HRP conjugate (catalog number: 4380-096-04, diluted 1:500-fold in assay diluent) was added to all wells, and incubated at 25 for 1 hour. The plate was then washed with 4 changes of 1xPBST with excessive liquid aspirated. 50 μL of room-temperature TACS-Sapphire colorimetric substrate (catalog number: 4822-096-08) was added to all wells and incubated for 15 minutes at 25 in the dark. The reactions were stopped by adding 50 μL of 5% phosphoric acid to all wells, and absorbance was read immediately at 450 nm. The average absorbance for all standards, samples and blanks was calculated, with the blank average subtracted from standard and sample averages to determine relative absorbance. The log of 8-oxo-dG standard concentration (ng/mL) was plotted against the relative absorbance to derive a standard curve, and 8-oxo-dG concentration in samples was calculated and interpolated from the standard curve.

### Terminal deoxynucleotidyl transferase dUTP nick end labeling (TUNEL) assay

Detection of apoptosis was carried out using ApopTag Fluorescein *In Situ* Apoptosis Detection Kit (S7110, EMD Millipore, MA, USA) following the protocol provided by the manufacturer. Briefly, slides were deparaffinized by passage through two changes of xylene (10 minutes each wash) and a descending series of ethanol (100%, 95%, and 70% for 5 minutes each wash), followed by one change of 1xPBS for 5 minutes. Slides were incubated in freshly diluted proteinase K (20 μg/mL) for 15 minutes at room temperature washed in 2 changes of 1xPBS for 2 minutes each wash. Equilibration buffer was added to each specimen and incubated at room temperature for 30 minutes. Immediately afterward, 55 uL of terminal deoxynucleotidyl transferase (catalog number: 90418, EMD Millipore, MA, USA) and reaction buffer (catalog number: 90417, EMD Millipore, MA, USA) mixture in 1:3 ratio was added to slides and stored in humidified chambers at 37 for 1 hour. Negative and positive controls were established by omitting the reaction buffer in the mixture and treatment with DNase I (catalog number: 4380-096-06, 5U/uL, Trevigan, MD, USA) at 37 for 1 hour, respectively. Reaction was stopped by incubating in Stop Buffer (catalog number: 90419, EMD Millipore, MA, USA) at room temperature for 10 minutes, followed by 3 washes in 1xPBS for 1 minute each wash. Room-temperature mixture containing anti-digoxigenin conjugate (catalog number: 90426, EMD Millipore, MA, USA) and blocking solution (catalog number: 90425, EMD Millipore, MA, USA) was added to each specimen and incubated at room temperature for 30 minutes in the dark. All slides were washed in 4 changes of 1xPBS for 2 minutes each wash, and counterstained with DAPI-containing ProLong Gold anti-fade mounting reagent. All specimens were stored at -20 in the dark. Images were captured on IX81 Olympus confocal microscope using the negative control as the minimal threshold for laser settings. All images were analyzed using ImageJ to manually count TUNEL-positive cells in each image (approximately 1000 cells per field), with the interfollicular epidermis and hair follicles quantified separately.

### Total RNA extraction from tissue and Next Generation Sequencing

RNA extraction from tissue was performed with TRIzol/chloroform extraction protocol. All equipment was cleaned with 70% ethanol and RNase-away solution (catalog number: 10328-011, Invitrogen, MA, USA). Briefly, approximately 1 gram of frozen skin biopsy and 500 μL of TRIzol (catalog number: 15596018, Life Technologies Corporation, CA, USA) were added into a micro-centrifuge tube and homogenized using a hand-held homogenizer. Fresh 500 uL of TRIzol were added to samples once they were adequately homogenized. After a 5 min incubation on ice, 200 μL chloroform (catalog number: 423555000, Acros Organics, NJ, USA) was added to samples. Samples were vortexed for 10 seconds and incubated at room temperature for 3 minutes. Samples were then centrifuged at 4 for 15 minutes at 12,000 rpm. The top aqueous layer was transferred into new tubes, ice-cold isopropanol (500 μL) was added, and samples were stored at -20 overnight. Samples were centrifuged at 4 for 30 minutes at 12,000 rpm, and supernatants were discarded. Pellets were washed in ice-cold 70% ethanol and centrifugation was repeated (4 for 10 min at 7,500 rpm). Pellets were air-dried and resuspended in 20 μL of nuclease-free ddH_2_O, with the RNA concentration measured via NanoDrop One/OneC (ThermoFisher Scientific, MA, USA). Samples with an A260/280 > 1.9 and A260/230 > 1.7 were used for further experiments and stored at -80 until further assays. RNA-sequencing was performed on RNA that had been isolated from six inbred *Hr^-/-^:Ptch1^+/-^*C57Bl/6 mice that had been UVB irradiated for 2 weeks. Half of the mice (n=3) were treated with HAPE; half of the mice (n=3) were treated with vehicle as a control. RNA-sequencing was performed at the University of Western Ontario Genomics Centre using the VAHTS Total RNA-seq (H-M-R) Library Prep Kit for Illumina with rRNA depletion from GeneBio Systems Inc using manufacturer’s instructions.

### Alignment of RNA-sequencing reads to GRCm39 reference genome

FASTQ files were imported into the Partek Flow (St. Louis, Missouri) bioinformatics platform according to manufacturer’s instructions, grouped into six samples, and assigned to one of two treatment groups: Vehicle Control or HAPE, depending on the type of topical cream treatment applied. Pre-alignment QA/QC was run on the samples to determine the quality of the unaligned reads. Each read underwent base trimming from the 3’ end, until bases with a Phred score of greater than or equal to 20 was reached. Trimmed reads were aligned to the GRCm39 reference genome using HISAT2 aligner (Version 2.1.0) with a whole genome aligner index.

### Differential expression analysis of aligned RNA-sequencing reads

Aligned RNA-sequencing reads were quantified to an annotation model (i.e., RefSeq Transcripts 98-2021-05-05 (Administrator)) using *Partek E/M* (according to manufacturer’s instructions) to quantify gene expression. Low expression genes were filtered out using a noise reduction filter (maximum ≤10). Read counts were then normalized to the total number of reads using the default settings. A value of 0.0001 was added to each gene’s normalized read count to eliminate mathematical errors resulting from values of zero. Differential expression analysis according to treatment group (i.e., HAPE or Vehicle Control) was then performed using *DESeq2*. Differential expression results were filtered by significance and fold change. The first round of filtering used thresholds of *p* ≤0.05, log_2_FC (fold-change) > 2 (−2>FC>2). The second round of filtering used thresholds of false discovery rate (FDR) ≤0.1, log_2_FC> 2 (−2>FC>2). Significantly differentially expressed genes according to these thresholds were then inputted into Metascape for gene ontology analysis^25^].

### Variant calling of RNA-sequencing data

Following alignment of RNA-sequencing reads to the reference genome, variant calling was performed using *SAMtools* (version 1.4.1). Variants were called against the GRCm39 reference genome, and variant annotation was performed using *RefSeq*. Variant data was then exported to Microsoft Excel, and all A>G variants were filtered out to account for RNA-editing artefacts ^26^]. Genes were then filtered according to whether they contained variants uniquely in HAPE-treated samples, uniquely in Vehicle Control-treated samples, or in both HAPE and Vehicle Control-treated samples. These unique gene lists were inputted into Metascape for gene ontology analysis^25^].

Following alignment of RNA-sequencing reads to the reference genome, variant calling was performed using *SAMtools* (version 1.4.1). Variants were called against the GRCm39 reference genome, and variant annotation was performed using *RefSeq*. Variant data was then exported to Microsoft Excel, and all A>G variants were filtered out to account for RNA-editing artefacts^26^] Genes were then filtered^26^] according to whether they contained variants uniquely in HAPE-treated samples, uniquely in Vehicle Control-treated samples, or in both HAPE and Vehicle Control-treated samples. These unique gene lists were inputted into Metascape for gene ontology analysis^25^].

### DNA extraction from tissue and whole exome sequencing

DNA extraction from tissue was performed using the Wizard Genomic DNA Purification Kit (Promega A1120) and following manufacturer’s instructions. WES was performed on DNA that had been isolated from four inbred *Hr^-/-^:Ptch1^+/-^* C57Bl/6 mice and that had been UVB irradiated for 26 weeks. Half of the mice (n=2) were treated with HAPE cream; half of the mice (n=2) were treated with Vehicle Control cream. WES was performed at the Centre for Applied Genomics Toronto using the SureSelectXT Mouse All Exon kit from Agilent according to manufacturer’s instructions.

### Alignment of whole exome sequencing reads to GRCm39 reference genome

WES FASTQ files were imported into the Partek Flow platform (used according to manufacturer’s instructions), grouped into four samples, and assigned to one of two treatment groups: Vehicle Control or HAPE, depending on the type of topical cream treatment applied. Pre-alignment QA/QC was run on the samples to determine the quality of the unaligned reads. Each read underwent adapter trimming using *Cutadapt* with the following adapters: 5’-CTGTCTCTTGATCACA-3’and 5’-TGTGATCAAGAGACAG-3’ (http://code.google.com/p/cutadapt/). Additionally, base trimming was performed from the 3’ end, until bases with a Phred score of greater than or equal to 20 was reached. Trimmed reads were aligned to the GRCm39 reference genome using *HISAT2* aligner (Version 2.1.0) with a whole genome aligner index.

### Variant calling of whole exome sequencing data

Following alignment of WES reads to the reference genome, variant calling was performed using *SAMtools* (version 1.4.1). Variants were called against the GRCm39 reference genome, variant annotation was performed using *RefSeq*, and variant data was exported to Microsoft Excel. Genes were then filtered according to whether they contained variants uniquely in HAPE-treated samples, uniquely in Vehicle-treated samples, or in both HAPE and Vehicle-treated samples. These unique gene lists were inputted into Metascape for gene ontology analysis^25^].

### Mutational signature analysis of RNA-sequencing and WES data

Code generated in-house was used to generate single base substitution (SBS) and double base substitution (DBS) mutational signature plots for each RNA-sequencing and WES sample. Cosine similarity matrices were generated to compare each mutational signature to one another, and to mutational signatures of interest from the COSMIC website^27^]. All WES and RNA-sequencing SBS mutational plots were compared to the following mutational signatures of interest: SBS1, SBS2, SBS5, SBS7(a-d), SBS17(a,b), SBS18, SBS38, SBS45, SBS46, SBS54, and SBS87. All WES and RNA-sequencing DBS mutational plots were compared to the following mutational signatures of interest: DBS1, DBS5, and DBS11.

### Spatial transcriptomics

To identify tissue blocks with intact RNA, ribbons of tissue sections were cut using RNAase free conditions and RNA was isolated using the RNAeasy FFPE RNA extraction kit (catalog number: 73504, Qiagen, ON, Canada) and following the manufacturer’s instructions. RNA concentration and size distribution was determined using the ABI 3730xI DNA Analyzer. RNA preparations with an RNA integrity number (RIN) of 2.4 and >30% of RNA fragments longer than 200 nucleotides (DV200) were considered to be of sufficient quality for sequencing. Sections of tissue blocks with sufficiently intact RNA were mounted on 10x visium slides (10x Genomics, CA, USA), stained for KRT15 and processed for RNA sequencing. Tissue sectioning, RNA quality analysis, processing of Visium slides, and RNA sequencing were performed at the London Regional Genomics Center, Western University. Sequencing data were processed using Space Ranger and Loupe browser 8 software (10x Genomics, CA, USA).

### Knockdown of TP53 expression in HaCaT cells

Cells were cultured as described above. 24 hours after plating, cells were either transfected with TP53ShRNA or treated with high molecular weight HA (1 mg/mL). 48 to 72 hours later, cells were fixed with 3% paraformaldehyde pH 7.5 (10 minutes at room temperature), washed with PBS (5 minutes at room temperature), treated with 0.1% Triton X100/PBS (5 minutes at room temperature), washed with PBS, then blocked with 3% BSA/PBS (1 hour at room temperature). Primary antibodies (cleaved Caspase3, Krt14, Krt15, TP53, Ki67) were diluted in 1% BSA/PBS, added on top of the coverslips and incubated over night at 4°C in a humidified chamber. The following day, coverslips were washed 3x in PBS (10 minutes at room temperature). Secondary antibodies were diluted 1:200 in 1% BSA/PBS then added on top of the coverslips followed by a 1-hour incubation at room temperature protected from light. To remove the secondary antibodies, coverslips were washed 3 x in PBS, 10 minutes each at room temperature. Coverslips were removed from 24 wells and mounted on microscope slides using ProLongGold antifade reagent mounting medium with DAPI (catalog number: P36931, ThermoFisher, OR, USA). Slides were imaged by confocal microscopy (IX81 Olympus confocal microscope equipped with FV10-ASW 4.2 software) and images were analyzed using ImageJ.

### HaCaT xenografts

6-week-old Nude (Nu/J) female mice were purchased from Jackson Laboratories (ME, USA). Housing, UV irradiation, subcutaneous injection, and euthanasia followed the approved animal protocol #2022-0117.

In brief, mice were exposed to 180 mJ/cm^2^ UVB on two days/week for 4 weeks as described above. HaCaT cells that were cultured in medium containing 1 mg/mL high molecular weight HA for 3 weeks were trypsinized, pelleted by centrifugation and mixed with Matrigel at a concentration of 2x10^7^ cells/mL. Following UVB irradiation, mice were anesthetized using Isoflurane inhalation and 50 μL of the Matrigel/cell mixture were injected into the right flank of Nude mice using a 31G needle attached to a 1 mL syringe. Tumor growth was monitored by palpation and caliper measurement twice/week for three weeks. Mice were euthanized by isoflurane inhalation followed by CO_2_ inhalation. Tumours were removed and fixed in 3% paraformaldehyde/PBS pH 7.5 and processed for histology. Immunofluorescent staining of tissue sections was performed as described above and images were captured either with a scanning confocal (IX81 Olympus) or a spinning disk confocal (Nikon CSUW1).

**Table 3:** Antibodies and buffers used in immunohistochemistry and immunofluorescence of paraffin-embedded tissue sections.

| Primary antibody | Catalog number | Dilution | Wash buffer | Antigen retrieval |
|---|---|---|---|---|
| GLB1 | A11132, Life Technologies Corporation, CA, USA | 1:200 | 1xPBS | 10 mM Sodium citrate, pH 6.0 |
| GLI1 | ab151796, Abcam, MA, USA | 1:900 | 1xTBS |  |
| CD44 | 553130, BD Biosciences, ON, CA | 1:200 | 1xPBS |  |
| HMMR | ab124729, Abcam, MA, USA | 1:50 | 1xPBS |  |
| H2AX (Ser139) | 9718, Cell Signalling Technology, MA, USA | 1:150 | 1xTBS |  |
| KI67 | ab16667, Abcam, MA, USA | 1:100 | 1xPBS |  |
| NFκB1 | sc-114, Santa Cruz, Biotechnology | 1:100 | 1xTBS |  |
| Cleaved CASP3 (Asp175) | 9664, Cell Signalling Technology, MA, USA | 1:50 | 1xTBS |  |
| KRT14 | ab181595, Abcam, MA, US | 1:250 | 1xPBS |  |
| KI67 (anti human) | ab92742, Abcam, MA, USA | 1:500 | 1xPBS |  |
| TP53 (anti human) | ab32049, Abcam, MA, USA | 1:500 | 1xPBS |  |
| Trp53 | ab1101, Abcam, MA, USA | 1:500 | 1xPBS |  |
| HA binding probe | K1200 Echelon, UT, USA | undiluted | 1xPBS | none |
| TUNEL assay | ab206386, Abcam, MA, USA | Manufacturer’s protocol |  |  |

## RESULTS

### HAPE penetrates the epidermis to form hyaluronidase-resistant coats around keratinocytes and block UVB-induced keratinocyte tumor initiation

Since topical HMW-HA does not enter skin due to its hydrophilicity^23^], we developed a co-polymer by linking HMW-HA (500 kDa) to phosphatidylethanolamine (HAPE) that readily penetrates skin. HAPE has been reported to form viscous coats around skin keratinocytes and fibroblasts in a receptor-independent manner^23^]. HAPE was applied topically to *Hr*^-/-^:*Ptch1*^LacZ/+^ mouse skin according to the schedule shown in Figure 1. HAPE is confirmed to penetrate the epidermis, form coats around keratinocytes to significantly increase epidermal HA retention compared to controls (Figure 2A) and suppress the expression of *Hyal2*, a common skin hyaluronidase, to the levels of unexposed skin (Figure 2B). HAPE also blocks the ability of hyaluronidase to remove HA coats on HaCaT keratinocytes in an *in vitro* particle exclusion assay (Figure 2C). Collectively, these properties mimic those of HMW-HA produced in the skin of naked mole rats^4,28^]. We next assessed if HAPE alters tumor initiation in response to UVB. Histology analyses show no difference in UVB-induced epidermal hyperplasia or number of benign lesions (papillomas) (Figure 2D). However, in contrast to vehicle controls, HAPE-treated mice do not develop keratinocyte tumors within the experimental time frame of 26 weeks (Figure 2D). The presence of *LacZ* on a *Ptch1* promoter allows tracking the activation of the main oncogenic driver pathway, SHH, for cutaneous basal cell carcinoma. The ability of HAPE to strongly suppress expression of one of the pathway target genes, *Ptch1*, provides molecular verification for suppression of the basal cell carcinoma initiation (Figure 2E).

**Figure 1.**
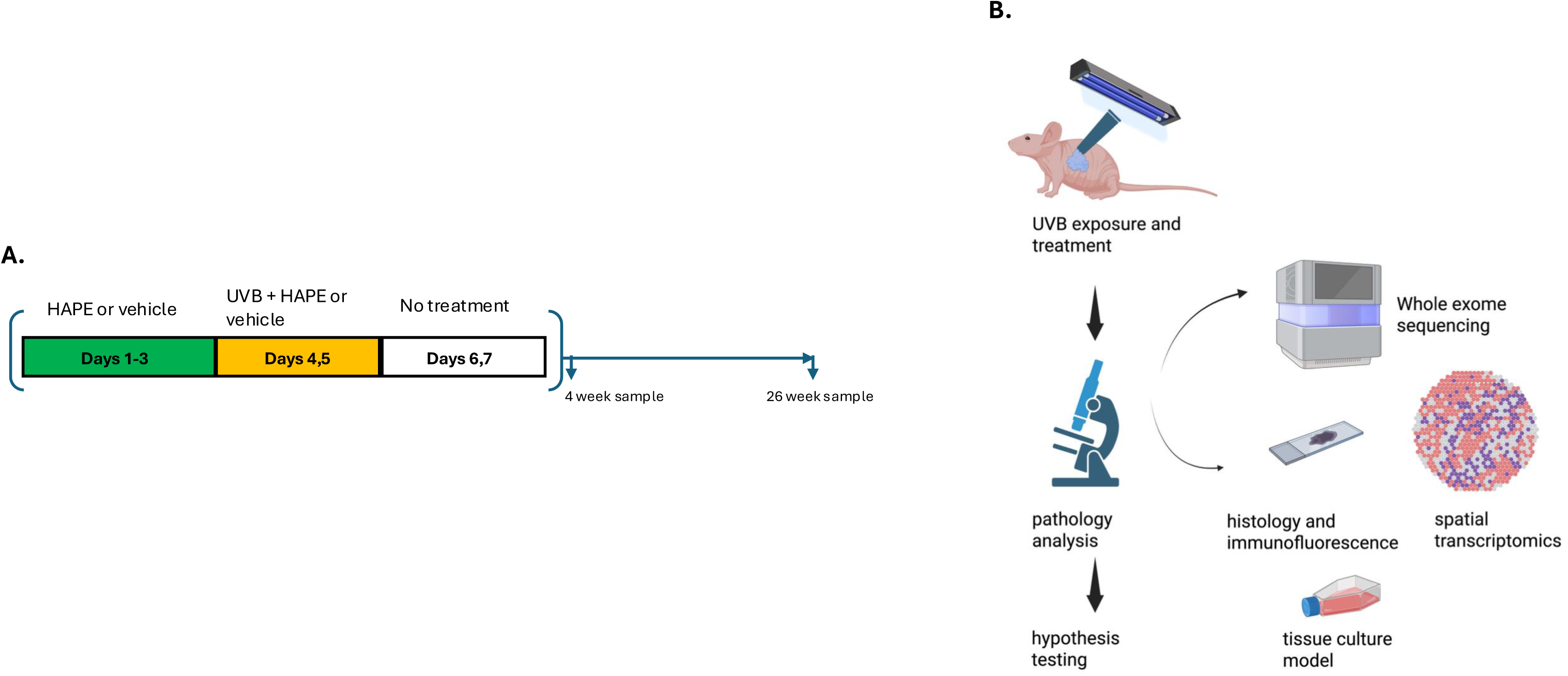
Schematic of experimental design. **A.** Experimental design and schedule for UVB exposure and treatment with HAPE, vehicle control or UVB alone. **B.** Treatment and analysis path for assessing tumorigenesis resulting from UVB exposure and tumor suppression by HAPE.

**Figure 2.**
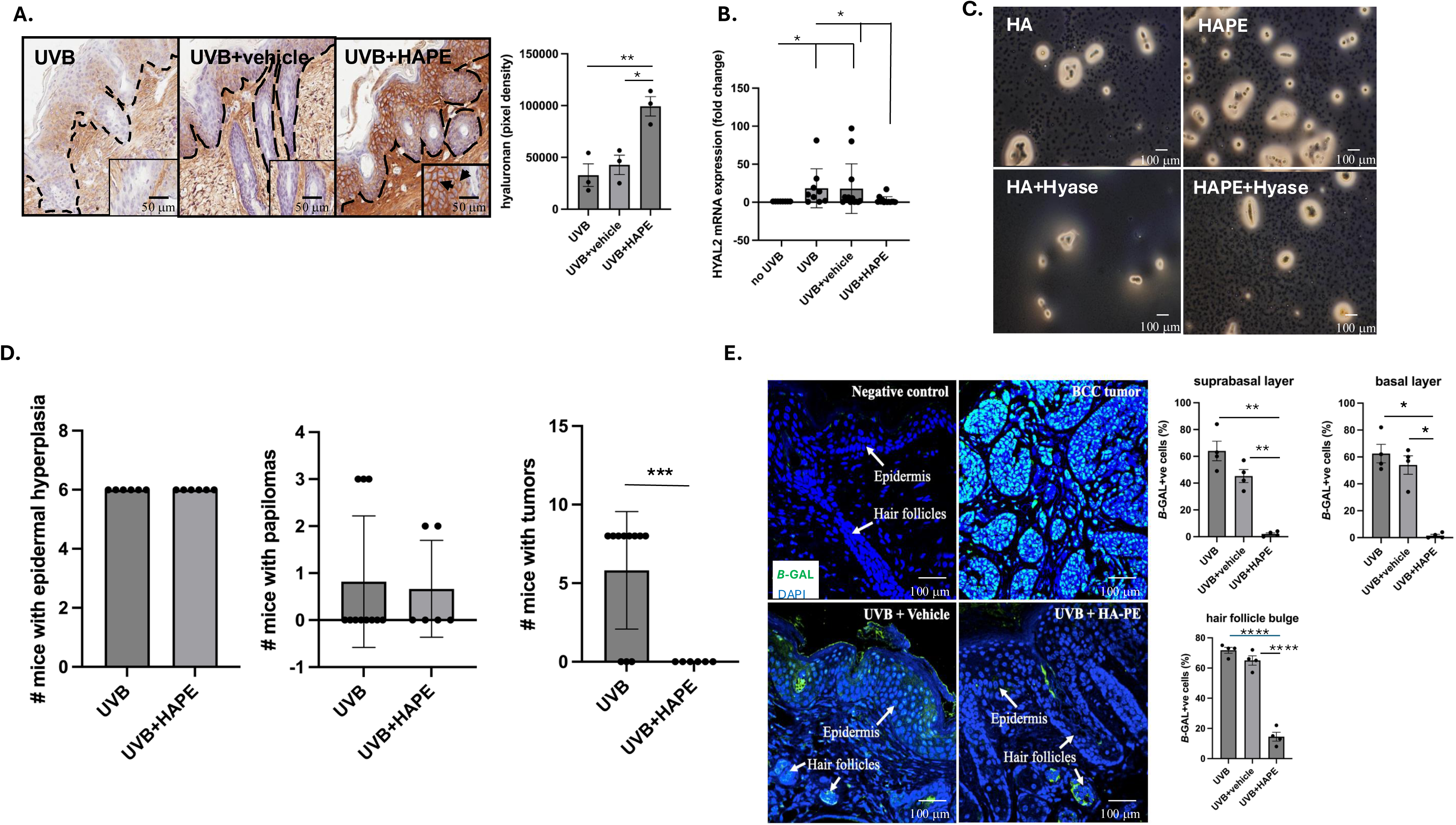
HAPE penetrates skin to form hyaluronidase-resistant coats around keratinocytes, which prevents keratinocyte tumor initiation. A. HAPE application increases HA staining in the epidermis and dermis vs. UVB and UVB + vehicle comparators. Inset shows that HAPE promotes the formation of HA coats shown to outline keratinocytes (arrow). B. UVB exposure increases hyaluronidase expression (HYAL2) in skin but HAPE application reduces expression of this enzyme. Values for A and B are the Mean and S.E.M. of N=3 biological replicates and N=6 technical replicates, *p<0.05, **p<0.01. C. Phase contrast microscopy of cultured HaCaT keratinocytes treated with either unmodified high molecular weight HA or HAPE using a particle exclusion assay. Micrographs show HAPE forming cell coats that are more resistant to hyaluronidase than unmodified high molecular weight HA. D. Pathology analyses of tissue sections show that HAPE does not prevent UVB-induced epidermal hyperplasia or benign papilloma formation but blocks appearance of cutaneous squamous and basal cell carcinomas. Values are the Mean and SEM. N=6 mice. ***p<0.001. E. Inhibition of cutaneous basal cell carcinoma initiation by HAPE is confirmed by quantifying B-GAL staining resulting from expression of LacZ protein driven by the *Ptch1* promoter. B-GAL staining is used as an indicator for the activation of SHH signaling, which is the oncogenic driver of basal cell carcinoma tumorigenesis. DAPI is used as the landmark nuclear stain. Negative control is stained with non-immune IgG. HAPE is shown to suppress oncogenic signaling in the suprabasal, basal and hair follicle bulge regions of skin. N=3 biological replicates and 6 technical replicates. Values are the Mean and SEM N=4 fields.

### HAPE promotes retention of basal:suprabasal polarity in the interfollicular epidermis and regulates proliferation and inflammation

Although the interfollicular epidermis (IFE) has undergone hyperplasia in HAPE-treated skin, epidermal polarity is retained as detected histologically by distinct basal and suprabasal layers (Figure 3A, B). Keratinocyte proliferation is also restricted to the basal layer of the IEF as is typical of skin that is not exposed to UVB (Figure 3B). Overall, HAPE reduces keratinocyte UVB-induced proliferation in the IEF and hair follicles (Figure 3B).

**Figure 3.**
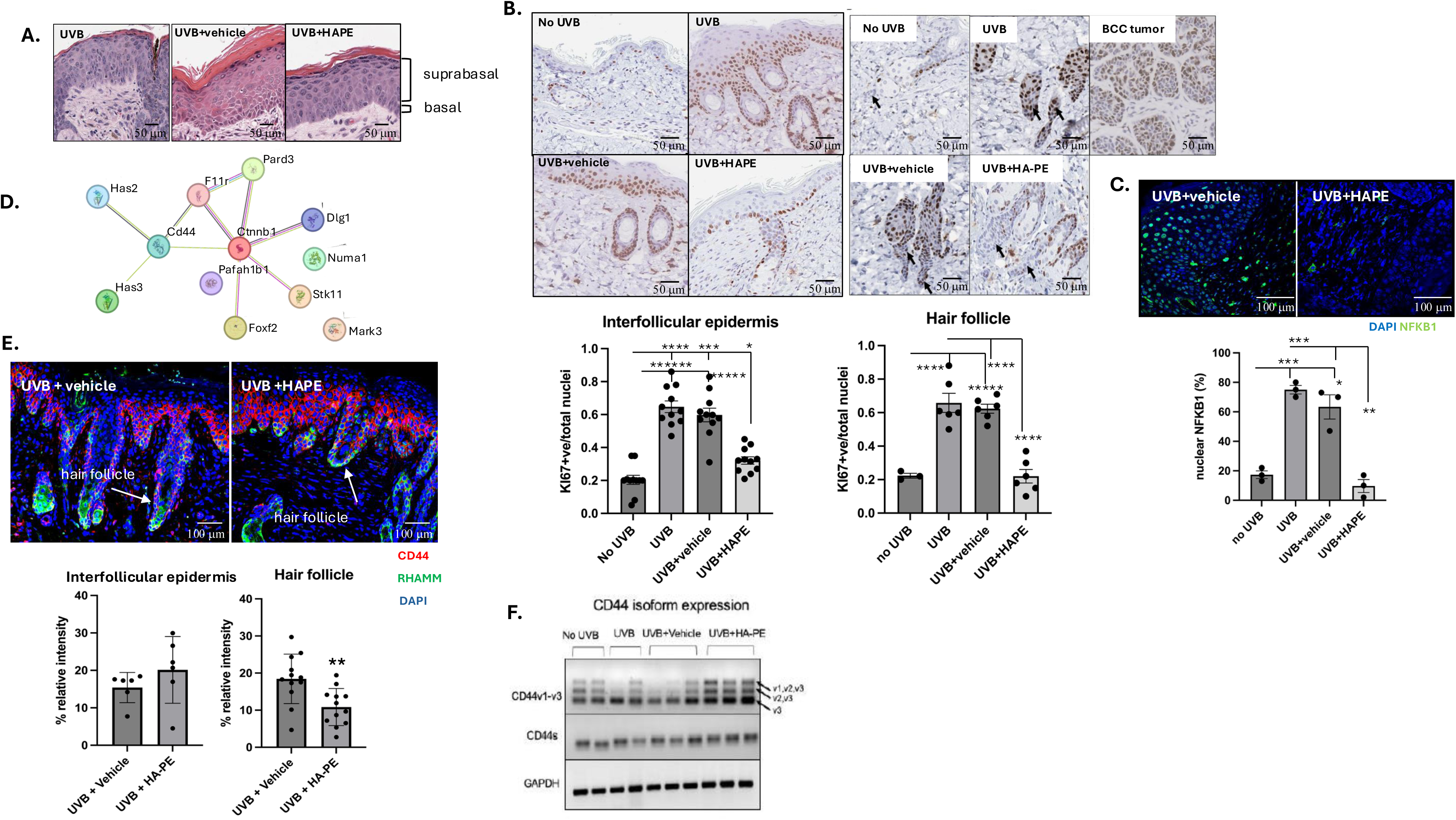
HAPE maintains epidermal quiescence in *Hr*^-/-^:*Ptch1^lacZ^*^/+^ mice. **A**. H&E paraffin-processed tissue section of UVB-exposed skin shows the organization of the epidermis into clearly discernable basal and suprabasal regions of skin with UVB+HAPE treatment. In contrast, the epidermis of UVB+vehicle and UVB alone controls is disorganized with loss of basal:suprabasal polarity. **B.** Cell proliferation, detected by KI67 staining, is increased in the interfollicular and follicular epidermis with UVB or UVB+vehicle treatment compared to unexposed skin. In contrast to UVB and UVB+vehicle controls, keratinocyte proliferation is not detected in the suprabasal layer of HAPE-treated interfollicular epidermis. Proliferation is elevated in the basal layer with UVB+HAPE treatment compared to unexposed skin but is less than UVB controls. UVB-induced cell proliferation in hair follicles is strongly reduced by HAPE to levels of unexposed skin. **C.** HAPE blunts inflammation as detected by reduced nuclear localization of NFKB1 (p50). **D.** STRING functional protein association network analyses predicts a connection between CD44/ hyaluronan synthases (HAS2,3) and proteins contributing to epidermal polarity (PPI enrichment p value=4.12e-05). Bulk RNAseq analyses confirms an HAPE-induced increase (p<0.05) in the expression of epidermal polarity marker genes relative to vehicle controls in UVB-exposed skin. **E.** Immunofluorescence shows that CD44 protein expression is slightly increased in the interfollicular epidermis but significantly decreased in hair follicles. **F.** Q-PCR shows that UVB and UVB+vehicle suppress keratinocyte-specific CD44 isoform mRNA expression normally detected in homeostatic skin. Expression of these isoforms is maintained with HAPE treatment. Values for **B**, **C**, and **D** are the Mean and S.E.M. *p<0.05, ** p<0.01, ***p<0.001, ****p<0.0001, *****p<1.0e-5, ******p<1.0e-6.

HMW-HA has been reported to have anti-inflammatory functions^29^] and inflammation has been shown to promote tumor initiation^30^]. We therefore also assessed if HAPE alters the expression and activation of NFKB1, detected by nuclear localization, which regulates pro-inflammatory target genes^31^]. While UVB exposure increases nuclear localization of NFKB1 in both the IEF and hair follicles, HAPE reduces this to levels of skin that has not been exposed to UVB (Figure 3C). Consistent with these results, bulk RNA-seq of skin shows a significant (p<0.05) increase in mRNA expression of polarity markers in HAPE-treated, UVB-exposed skin. Furthermore, STRING functional protein association network analyses of these markers predict an association of HA metabolism, in particular the expression of the HA receptor, CD44 to expression of epidermal polarity genes (Figure 3D). Since CD44 is linked to the tumor suppressive effects of

Naked mole rat (NMR-HA^1^]), we quantified its expression in HAPE vs. vehicle-treated UVB-exposed skin. Immunofluorescence analysis detected similar levels of CD44 in the interfollicular epidermis (IFE), HAPE downregulated CD44 protein in hair follicles (Figure 3E). Isoform specific PCR analyses (Figure 3F) show that CD44 mRNA expression is significantly upregulated by HAPE and that the keratinocyte specific CD44 isoform (V1-3) normally expressed by skin is reduced by UVB exposure and that HAPE restores V1-3 expression (Figure 3F). Collectively these results show that HAPE preserves the skin polarity and homeostasis that is lost in *Hr*^-/-^:*Ptch1*^LacZ/+^ mice exposed to UVB.

### HAPE does not prevent UVB-induced DNA damage and mutations in oncogenes

To begin to dissect the mechanisms by which HAPE preserves epidermal polarity, its effect on UVB-induced DNA damage and/or mutations was next assessed. Early DNA damage was detected using the biomarker phospho^Ser139^-gamma-H2AX (Figure 4A). Similarly extensive nuclear staining was observed throughout the epidermal layers of HAPE and vehicle control in the IFE and hair follicles of UVB-exposed samples. Bulk UVB-induced DNA damage was further quantified by measuring 8-Oxo-dG levels resulting from elevated reactive oxygen species and direct UVB photon absorption^32^]. The levels of this damage are strongly increased in UVB vs. unexposed skin but are not significantly different in HAPE vs. vehicle control samples (Figure 4B).

**FIGURE 4.**
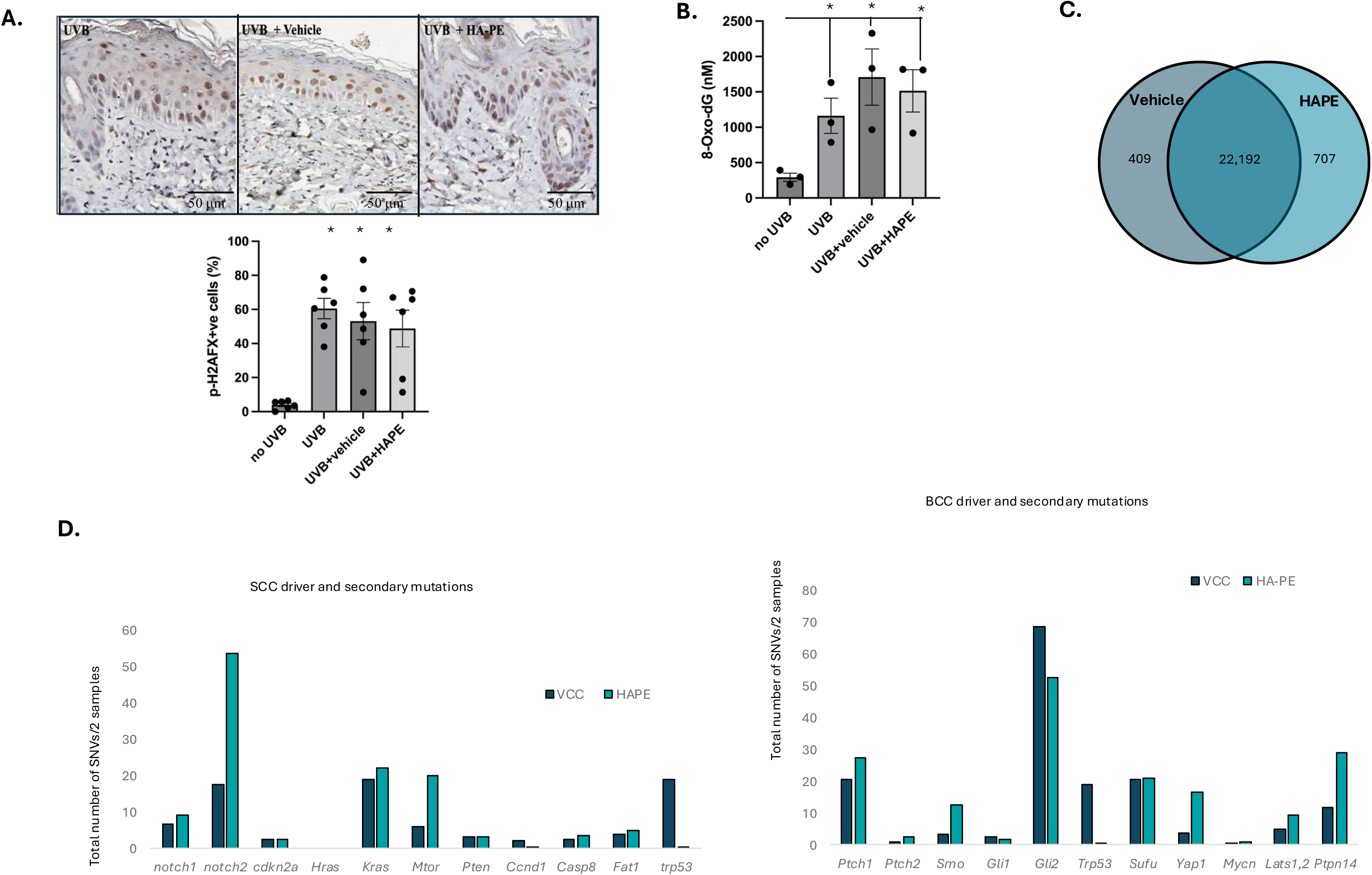
UVB-induced DNA damage and mutagenesis are unaffected by HAPE. **A.** Early DNA damage is detected by double-strand breaks marked by phospho-H2AX staining. UVB increases staining in vehicle and HAPE treated skin. **B.** UVB directly damages DNA and promotes reactive oxygen species-induced DNA damage, detected by 8-oxo-dG levels. HAPE does not prevent this UVB-mediated DNA damage relative to vehicle-treated UVB controls. **C.** Whole exome sequencing (WES) of DNA detects a similar total mutation burden (single nucleotide variants, SNVs) in control (vehicle) and HAPE-treated skin exposed to UVB. Most of the mutations detected by WES are shared in vehicle control and HAPE-treated, UVB-exposed skin Data are shown as a Venn diagram. **D.** Analyses of WES identify SNV mutations in oncogenic driver genes for cutaneous squamous cell (SCC) and basal cell (BCC) carcinomas. Mutations in these oncogenic driver genes are similar in vehicle control and HAPE-treated skin except for *Trp53*, which is not detected in UVB-exposed HAPE treated skin. Values for **A**, **B** are the Mean and S.E.M. *p<0.05, **p<0.01. WES analyses are from two biological replicates.

We next assessed if the DNA damage detected in HAPE-treated samples results in a mutation burden. Since the DNA damage resulting from UVB exposure primarily causes single nucleotide variants (SNVs) that drive cancer initiation and evolution^33^], we performed whole exome sequencing (WES), which detects SNVs and indels^34^]. WES analysis shows that SNVs are present in both UVB-exposed vehicle control- and HAPE-treated skin, and the majority of SNVs are shared between control and HAPE-treated skin (Venn Diagram, Figure 4C). Querying variants using the COSMIC database show that both bear similar UV mutation signatures (S-Figure 1). The mutation load of specific oncogenic driver, suppressor and secondary genes for cutaneous squamous and basal cell carcinomas was next documented (Figure 4D). The pattern of SNVs in oncogenic driver genes is similar in control and HAPE-treated skin with the notable exception of *Trp53.* Only 1 SNV in this gene is detected in one sample of HAPE-treated skin compared to an average of 20 SNVs in each vehicle control sample. Overall, these results show that a UVB-induced cancerized field of mutations is established in both HAPE and control skin. One prediction deduced from these results is that HAPE acts to suppress the function of oncogenic mutations, which blocks keratinocyte tumor initiation.

### HAPE reduces expression of SHH pathway targets and TRP53

We first considered the possibility that HAPE suppresses expression of mutant oncogenes required for cancer initiation and present in the cancerized field. Bulk RNA-seq of skin punches was performed as a high-level unbiased initial screen of differentially expressed genes but targets primarily affected muscle function likely due to the panniculus carnosus muscle located beneath the dermis of mouse skin preventing assessment of oncogenic signaling. We therefore quantified the expression of oncogenic pathway targets using immunofluorescence of histology tissue sections. This allows analyses of each epidermal compartment particularly within the interfollicular epidermis and hair follicles where cancer initiating progenitor cells reside^20^]. We queried the effect of HAPE on genes known to be involved in keratinocyte tumor initiation specifically SHH, required for cutaneous BCC initiation and TRP53, which is regulated by NMR-HA^35^] and whose functional loss contributes to escape of cutaneous BCC from dormancy and to SCC progression^36,37^], and enhances cBCC carcinogenesis in *Ptch1*^+/-^ mice^21^] and other experimental models^38^].

Elevated expression of GLI1 and SHH were used as evidence for the activation of the SHH pathway, an oncogenic driver of cBCC. As expected, GLI1 is strongly expressed in BCC tumors and in the IEF and hair follicles of UVB+vehicle (hair follicles, insert, Figure 5A). HAPE suppresses expression of GLI1 to levels observed in samples that are not exposed to UVB (Figure 5A). SHH mRNA expression is also increased in the UVB+vehicle control and suppressed by HAPE to levels of skin not exposed to UVB (Figure 5A). Immunofluorescent analyses further show that SHH pathway activation by UVB and its suppression by HAPE is widespread affecting IFE and hair follicles.

**FIGURE 5.**
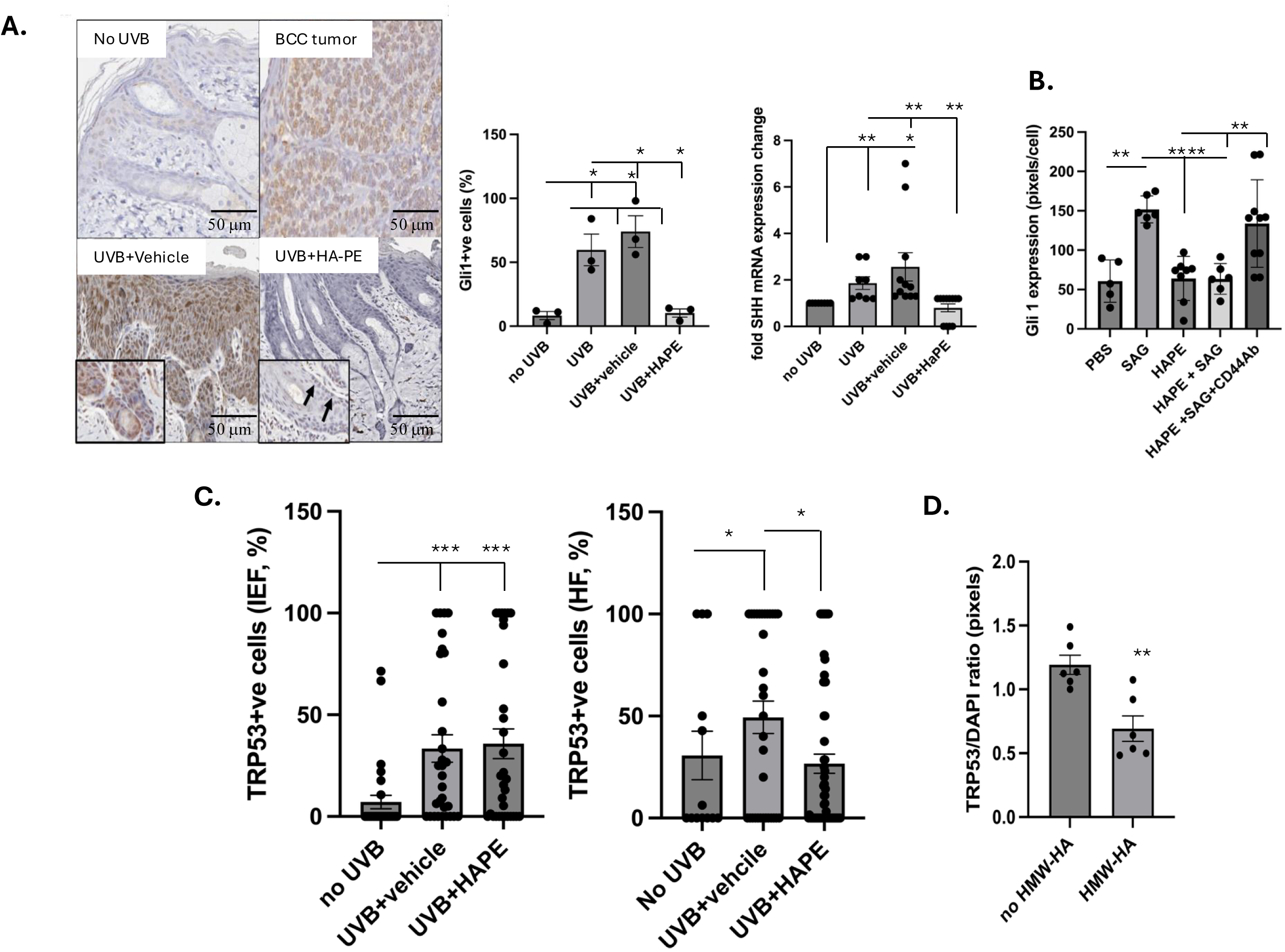
HAPE suppresses oncogenic signaling. **A.** HAPE strongly suppresses GLI1 and SHH expression. GLI1 protein expression was detected by immunohistochemistry and quantified using Image J while SHH mRNA expression was quantified by qPCR. **B**. In cultured human (HaCaT) keratinocytes, expression of GLI1 requires CD44 signaling, which HAPE suppresses. **C**. HAPE suppresses TRP53 protein expression in hair follicles (HF) but not in the interfollicular epidermis (IEF). **D.** HAPE suppresses protein expression TRP53 in HaCaT keratinocytes. Values are the Mean and S.E.M. *p<0.05, **p<0.01, ***p<0.001, ****p<0.0001.

Experimental manipulation of the human HaCaT keratinocyte line confirms that activation of SHH signaling expression by a SMO agonist (SAG), detected by quantifying GLI1 protein expression, is blocked by HAPE (Figure 5B). CD44 expression is required for this effect since a function blocking anti-CD44 antibody abrogates this effect of HAPE (Figure 5B). These results are consistent with previous evidence that NMR-HA suppresses fibroblast transformation *in vitro* via engagement with CD44^1^].

In contrast to the SHH pathway, the effect of HAPE on TRP53 expression is compartmentalized to a specific area of the epidermis. UVB increases the expression of TRP53 protein in both the IFE and hair follicles above unexposed skin for the vehicle control and HAPE samples (Figure 5C). While UVB+HAPE also stimulates TRP53 protein expression in the IFE, it suppresses expression in hair follicles (Figure 5C). HMW-HA similarly reduces TRP53 protein expression in HaCaT keratinocytes *in vitro*, which contain KRT15+ cells, a cancer-initiating cell marker expressed in hair follicles and keratinocyte tumors (Figure 5D).

These results indicate HAPE strongly suppresses SHH signaling occurring throughout a UVB-induced cancerized field. In contrast to this widespread effect, HAPE-induced suppression of TRP53 expression is limited to hair follicles. The collective results suggest HAPE blocks keratinocyte tumorigenesis by at least two distinct mechanisms that target oncogenic SHH activity and TP53 expression/function. Since TRP53 function impacts both cutaneous SCC and BCC initiation/progression as well as cancer initiating cells^39^] and is regulated by HMW-HA^6,40],^ we probed the effects of HAPE on cancer-initiating cells residing in hair follicles.

### HAPE promotes the death of TRP53+:KRT15+ subsets in hair follicles

Under homeostatic conditions KRT14 is primarily expressed in progenitor cells of the IFE basal layer and in the outer root sheath of hair follicles while KRT15 expression is normally limited to the hair follicle bulge. Since both KRT14+ and KRT15+ stem cells can initiate keratinocyte tumors in experimental models exposed to UVB or gamma radiation^21,41–45^], we examined the effect of UVB with and without HAPE on the expression of these keratins. KRT14 and KRT15 expression occur throughout the IFE layers and in hair follicles with UVB exposure (Figure 6B). The treatment of UVB-exposed skin with HAPE reduces expression of both keratins in the IFE and hair follicles (Figure 6B). Since KRT15 hair follicle cells are specifically cancer-initiating in UVB-exposed *Hr*^-/-^*Ptch1*^LacZ/+^ mice^22^], we focused our analyses on these hair follicle progenitor cells. At 4 weeks of exposure to UVB, hair follicle cells express similar amounts of KRT15 in vehicle and HAPE-treated samples (Figure 6C). However, by 26 weeks of UVB exposure, KRT15 expression in hair follicles is significantly less in HAPE-treated vs. vehicle controls (Figure 6C). Loss is associated with the death of hair follicle KRT15+ subsets as detected by staining overlap with apoptosis markers, cleaved CASP3 (Figure 6D) and TUNEL (Figure 6E). KRT15+ loss also coincides with a reduction in KRT15+/TRP53+ cells and in TRP53 activity detected by staining for TRP^S15^ (Figure 6F, G). We next performed spatial transcriptomics on KRT15-stained histology slides to further characterize the subsets that HAPE may target.

**FIGURE 6.**
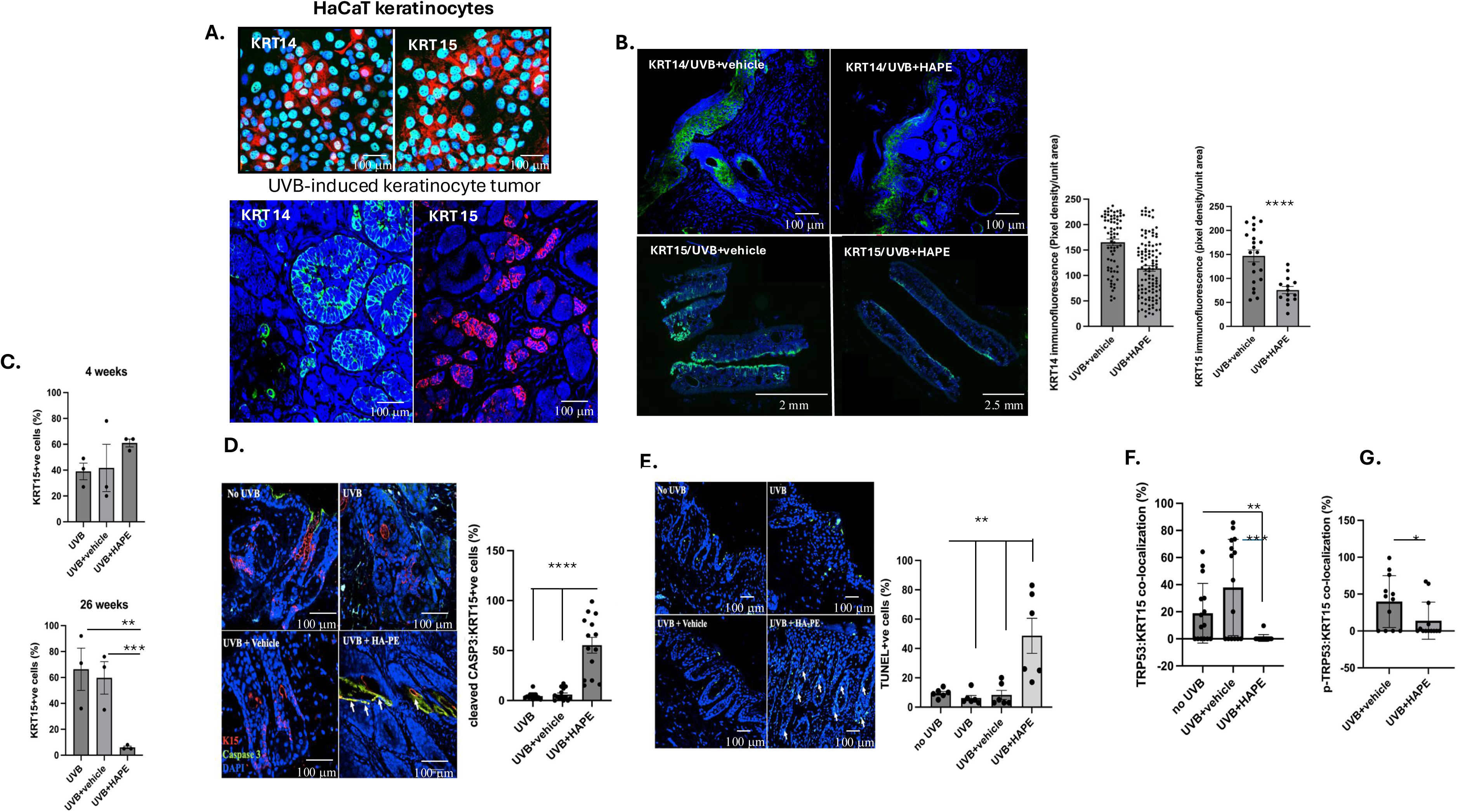
HAPE promotes apoptosis of KRT15+ hair follicle subsets. **A.** HaCaT keratinocytes and UVB-induced keratinocyte tumors express the keratinocyte stem cell markers, KRT14 and KRT15. **B.** The staining for KRT14 and KRT15 is significantly decreased in UVB-exposed skin of HAPE vs. vehicle controls as quantified by Image J. **C.** HAPE promotes the loss of KRT15+ve hair follicle progenitor subsets between 4 to 26 weeks of UVB exposure. **D, E.** Immunofluorescent analyses show that hair follicle KRT15+ stem cell subsets undergo apoptosis with HAPE treatment as detected by cleaved Caspase 3 (**D**) and TUNEL staining (**E**). **F, G.** HAPE reduces the expression (F) and activation (G) of TRP53 in KRT15+ve subsets. Values are the Mean and S.E.M. *p<0.05, **p<0.01, ***p<0.001, ****p<0.0001.

### Spatial transcriptomics identifies a KRT15+:TRP53+ hair follicle subset targeted by HAPE

We first identified differences between tumors and the surrounding histologically normal areas of vehicle controls that had developed field mutations due to UVB exposure. Keratinocyte tumors express higher levels of oncogenic driver and keratinocyte stem cell marker genes (Figure 7A, B) compared to the surrounding histologically normal epidermis. These oncogenic and stem cell marker genes are predicted to control keratinocyte fate and SHH signaling (Figure 7C). Pathway analyses (GSEA molecular signature data base, Hallmark gene sets, p<0.05) of genes upregulated in tumors vs. surrounding skin predicts altered tumor cell plasticity (EMT), increased oncogenic signaling, response to UV and modification of microenvironmental cues (e.g. angiogenesis) (Figure 7D). These results confirm that tumors are distinct from the surrounding UVB-exposed skin by expressing higher levels of oncogenic driver genes and progenitor markers. We next assessed the consequences of HAPE on transcriptomes of UVB-exposed healthy skin to detect differences that might account for the suppression of tumor initiation.

**FIGURE 7.**
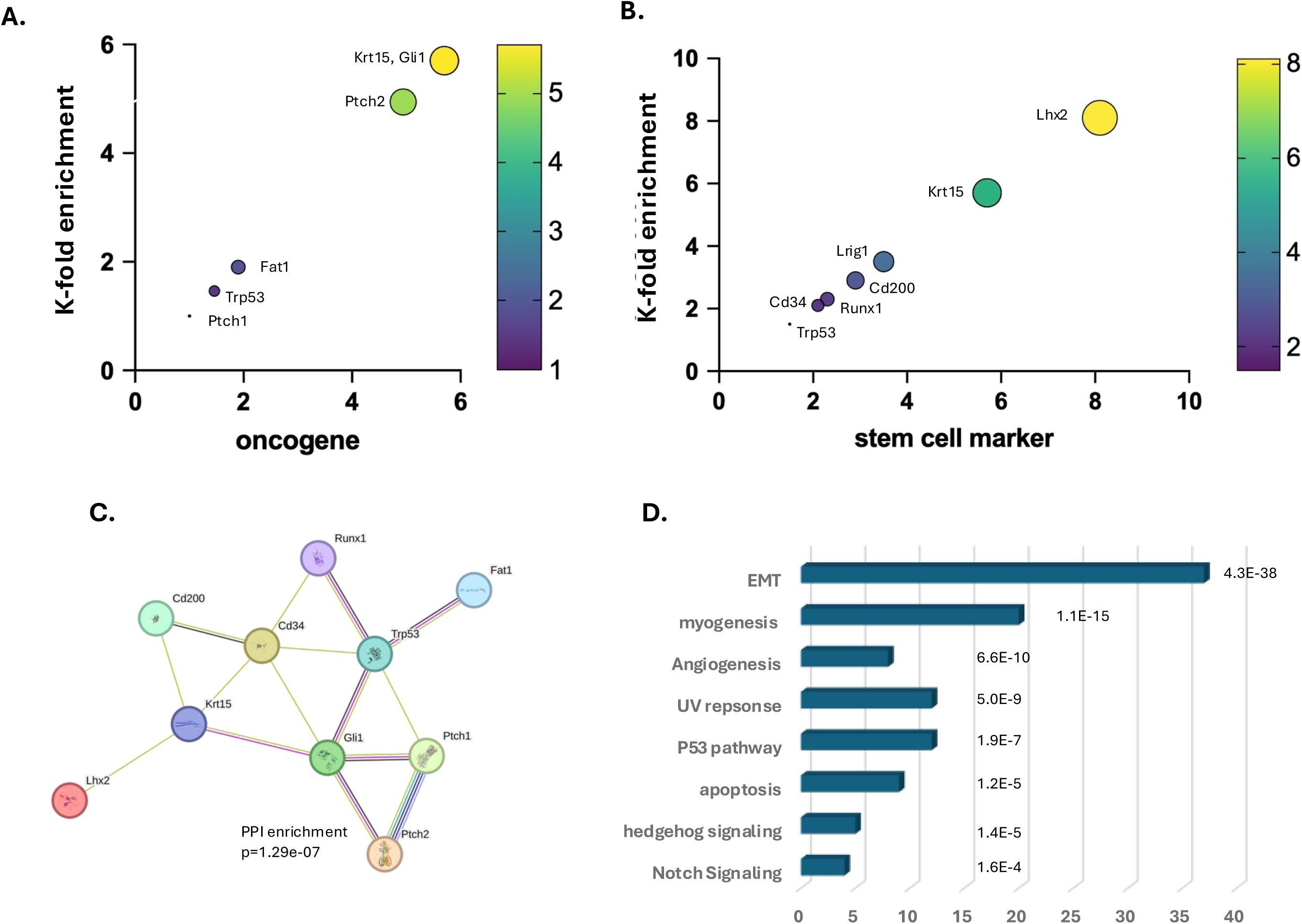
Spatial transcriptomic analyses show that oncogenic driver gene and stem cell biomarker expression are higher in tumors than the surrounding histologically normal skin. A., **B.** Dot blots showing the differentially expressed genes (DEG) of oncogenic driver genes (A) and stem cell markers (B) for tumors vs. surrounding, “cancerized skin”. C. Protein-protein interaction networks functional enrichment analysis (<u>string-db.org</u> ) predicts enrichment for epithelial development and epidermal fate specification as well as hedgehog signaling in tumors. D. Pathway enrichment using Gene Set Enrichment Analysis (GSEA), molecular signature data base (https://www.gsea-msigdb.org/gsea/index.jsp ) predicts significantly (p<0.05) upregulated oncogenic functions in tumors vs. surrounding skin in particular altered tumor cell plasticity (epithelial to mesenchymal transition, EMT) and increased oncogenic driver signaling.

UMAP images, built from 2 biological replicates and 4 technical replicates/treatment, identified 19 spatially distinct transcriptomic clusters using a 0.2 log fold change to identify differentially expressed genes that are present in HAPE and vehicle controls (Figure 8A). The analyses for vehicle controls used UVB-exposed skin that did not contain tumors and was matched to HAPE-treated samples for location. The boundaries between the clusters are more defined in HAPE vs. vehicle treated skin but both transcriptomes are typical for keratinocytes, hair follicles and myocytes in mouse skin (Figure 8A). Pathway analysis using GSEA datasets (canonical pathways) predicts that oncogenic signaling and response to UV exposure are upregulated in the vehicle controls vs. HAPE-treated UVB-exposed skin (Figure 8A). Dot plot analyses show that expression of cutaneous SCC and BCC oncogenes, HA metabolism and stem/progenitor cell biomarkers are elevated in vehicle controls compared to HAPE-treated, UVB-exposed skin (Figure 8B, C). These results confirm that HAPE blunts oncogenic signaling via blocking expression of pathway components.

**FIGURE 8.**
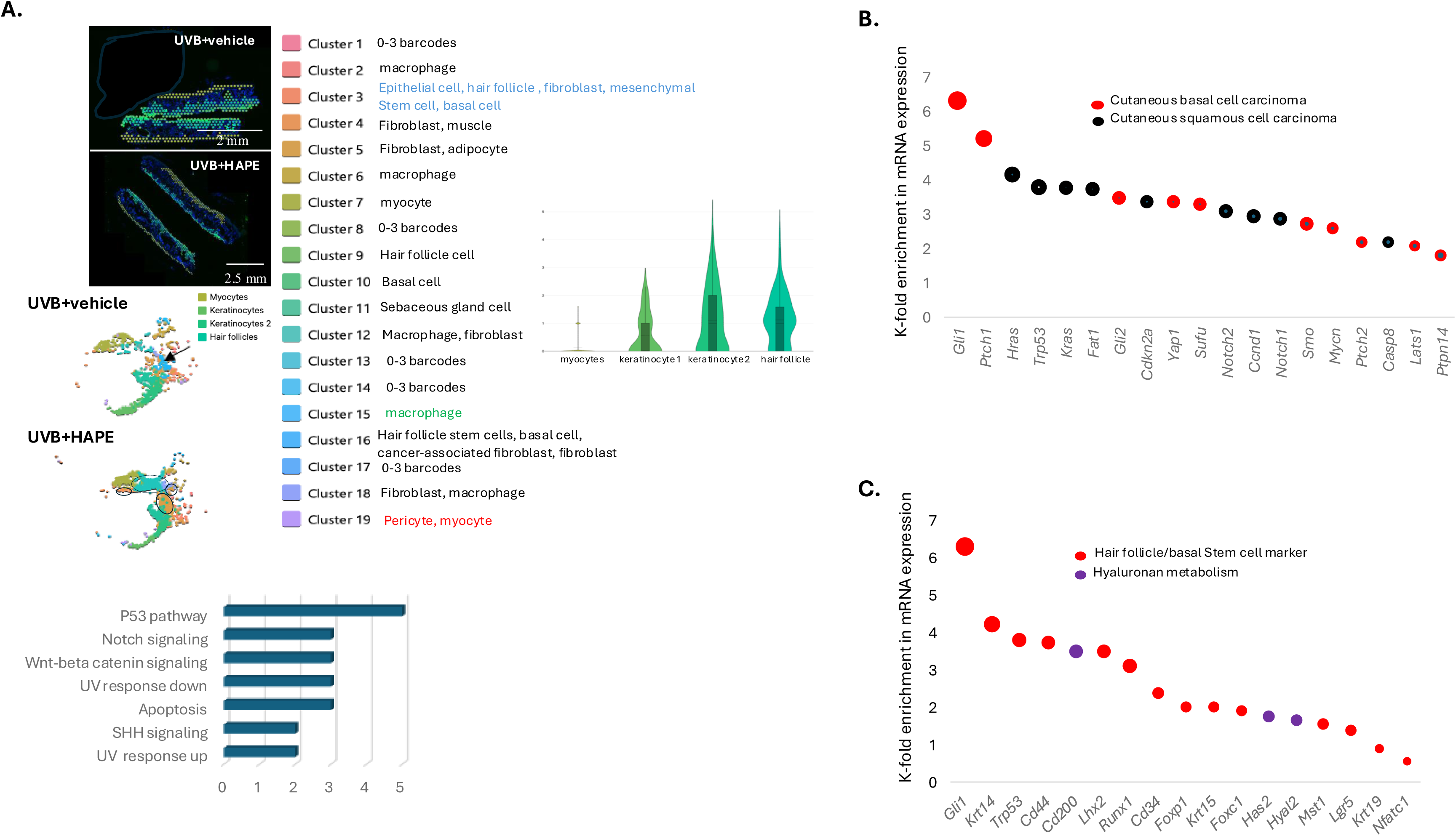
Spatial transcriptomic analyses of UVB-exposed skin tissue sections identify changes in oncogenic functions with HAPE treatment. Analyses confirm HAPE-mediated reduction in HA metabolism, hair follicle and basal stem cell marker expression and oncogenic pathway signaling. A. UMAP and cell annotation analyses were performed on the tissue sections stained with KRT15. The major cell types present in the clusters of UMAPs were identified using CellRanger (10X genomics). Violin plots from these analyses confirm that KRT15 expression is limited to keratinocytes and hair follicle cells. UMAP clustering is based on 19 principal components from HAPE and vehicle-treated skin using a 0.2 log fold change to identify differentially expressed genes and were built from N=2 biological replicates and 4 technical replicates/treatment. Each cluster corresponds to a distinct spatial location in skin sections mounted on visium slides. Pathway analysis using GSEA datasets (canonical pathways) predicts that oncogenic signaling and response to UV exposure is suppressed by HAPE. **B., C.** Dot plots show the HAPE-mediated decrease (p<0.05) in expression of individual stem cell markers (B) as well as HA metabolism and oncogenic driver genes (C) vs. UVB treated samples.

UVB-induced mutations in p53 are required for progression of cutaneous SCC^46^] and for escape of BCC from dormancy to macroscopic disease^37,47^]. Since WES analyses predict that *Trp53* mutations are rare in HAPE-treated UVB-exposed skin (Figure 4D), we hypothesized that a critical anti-tumor function of HAPE is to remove progenitor cells bearing mutant *Trp53*. We reasoned that critical KRT15+/mutant Trp53 subsets would therefore be lost with HAPE treatment. We therefore re-sorted spatial transcriptome data of differentially expressed genes into KRT15+/TRP53+ transcriptomic clusters (Figure 9A, B). Spatial transcriptomics analyses uncover 3 TRP53+ (arrows, Figure 9B) and one TRP53+/KRT15+ cluster (cluster 3, arrow, Figure 9A). All 3 clusters contain cells that express stem cell markers (Figure 9C) but only cluster 3 is predicted to contain hair follicle cells (Figures 8A, 9D). Importantly, TRP53 expression is lost, and KRT15 expression is reduced, in cluster 3 following HAPE treatment. GSEA pathway analyses predict EMT, K-RAS and TRP53 signaling downregulation, and upregulation of hair follicle differentiation pathways for this cluster (Figure 9D). 63% of the TRP53 target genes reported by 7 independent research groups^48^] are reduced by HAPE (Figure 9E) while 69% of the TRP53 target genes regulated by NMR-HA^40^] are altered by HAPE (Figure 9F), confirming TRP53 pathway shutdown as predicted by p-TRP53 immunofluorescence (Figure 6G). These results together with previous evidence that somatic mutations in TP53 of normal cells are generally not detected in other tissues^49^] prompted us to further analyze the consequence of HAPE on KRT15+ keratinocyte subsets bearing mutant TP53.

**FIGURE 9.**
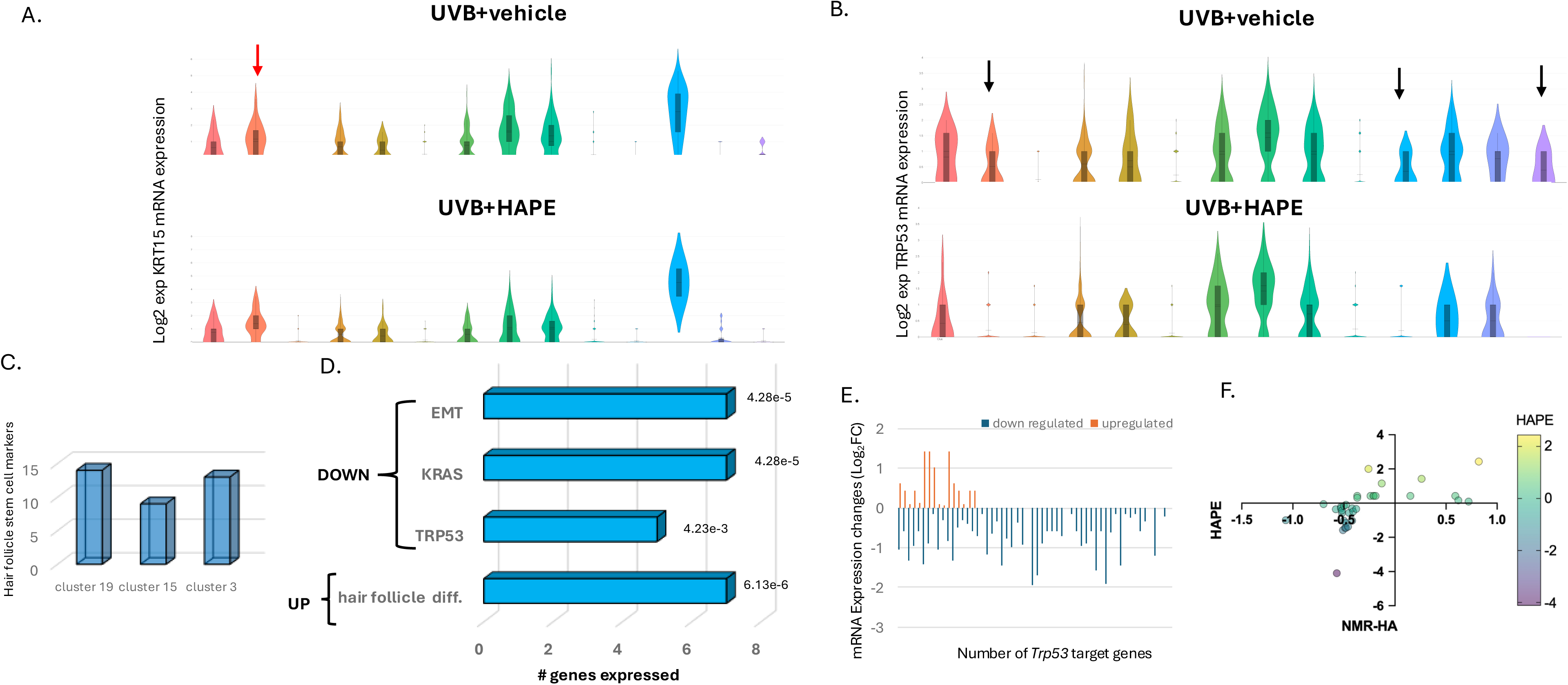
HAPE treatment causes the loss of distinct subsets. A, **B**. Violin plots of cell subsets from analyses shown in Figure 8A that were sorted for KRT15 (A) and TRP53 (B) mRNA expression. Three subsets (black arrows, **B**, subsets 3, 12, 19) expressing TRP53 mRNA in UVB+vehicle disappear in the HAPE-treated comparator one of which is KRT15+ve (red arrow, **A**, subset 3). **C.** All 3 HAPE depleted subsets express stem cell markers but cluster 3 and 19 exhibit the highest number of hair follicle stem cell markers. **D.** Functional annotation of HAPE-driven differentially expressed genes in cluster 3 predicts downregulated genes are involved in EMT, KRAS and TRP53 signaling while upregulated pathways promote hair follicle differentiation. **E.** TRP53 gene targets are largely downregulated in cluster 3 including those reported to be regulated by NMR-HMW-HA (**F**).

UVB results in both loss- and gain-of-function mutations in TRP53^50^]. Theoretically, reduction of TRP53 protein with loss-of-function mutations should not elicit apoptosis but reduction of TRP53 protein with gain-of-function mutations should promote apoptosis^51^]. We therefore hypothesized that HAPE promotes apoptosis of KRT15+ progenitors that acquire gain-of-function *Trp53* mutations. To test this hypothesis, we utilized the human keratinocyte line, HaCaT, that was isolated from a UV-exposed, cancerized field, contains gain-of-function mutations in TP53^52^] and undergoes apoptosis^51,53^] when expression of the mutant protein is suppressed. As described in Methods, Nude mice were exposed to UVB for 4 weeks to establish a cancerized field of oncogenic mutations prior to HaCaT cell injection.

We first confirmed that knockdown of TP53 expression by shRNA increases apoptosis of HaCaT keratinocytes (^51^], Figure 10A). We further showed that knockdown of TP53 promotes loss of a TP53+:KRT15+ subset of HaCaT keratinocytes (Figure 6A, 10B). Unmodified HMW-HA was added to cells *in vitro* and has no detectable effect on the number of KRT14+ HaCaT cells but reduces those of KRT15+ comparators increasing their apoptosis as detected by cleaved CASP3 staining (Figure 10D). Therefore, the ability of HMW-HA to target mutant TP53+/KRT15+ HaCaT subsets was next assessed. Immunofluorescence shows that HMW-HA promotes the loss of TP53+/KRT15+ HaCaT keratinocyte subsets (Figure 10E).

**Figure 10.**
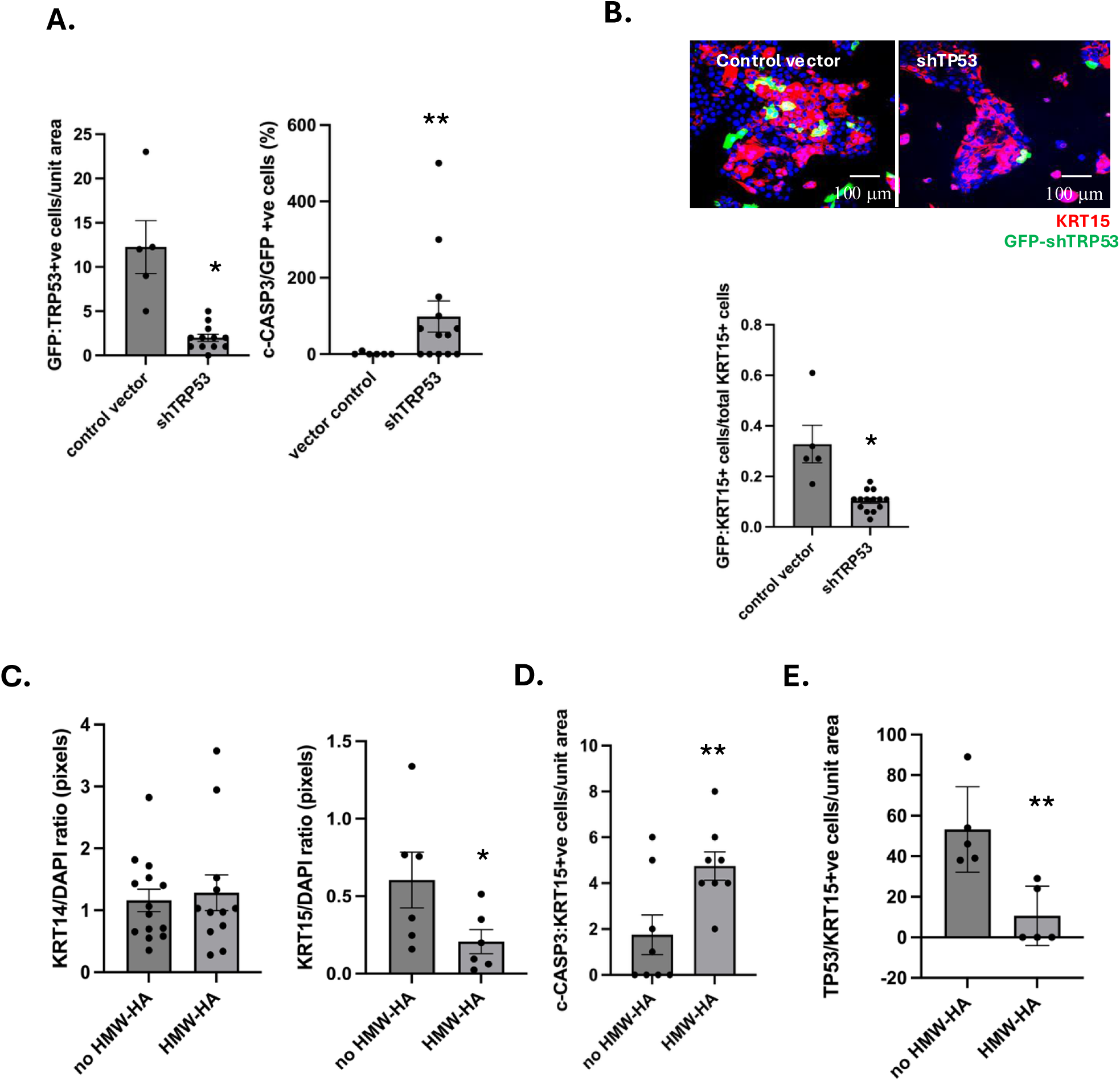
High molecular weight HA (HMW-HA) pretreatment increases apoptosis of TP53+:KRT15+ HaCaT subsets *in vitro*. **A.** Knockdown of TP53 (GFP-SHTP53) expression decreases HaCaT keratinocyte survival and increases apoptosis as indicated by cell loss and enhanced staining for cleaved CASP3. **B.** TP53 knockdown decreases survival of KRT15+ve HaCaT keratinocytes. **C.** HMW-HA promotes loss of KRT15+ve but not KRT14+ TP53+ HaCaT keratinocytes, quantified as a ratio of DAPI+ve adherent cells. **D**. HMW-HA increases apoptosis of KRT15+ve HaCaT keratinocytes as detected for immunofluorescence of cleaved CASP3. **E.** HMW-HA causes loss of TP53+/KRT15+ HaCaT keratinocytes. Values are the Mean and S.E.M. *p<0.05. **p<0.01, ***p<0.001.

HaCaT cells have been reported to form small, invasive tumors in Nude mice when they are injected subcutaneously as spheroids grown in Matrigel^54^]. We used this method to assess the effect of exposing HaCaT/Matrigel spheroids to HMW-HA on subsequent growth patterns as xenografts in Nude mice exposed to UVB to establish a cancerized field of mutations. HMW-HA-treated HaCaT/matrigel spheroids were injected subcutaneously on back skin. Histology of H&E-stained tissue sections shows that HaCaT keratinocytes form disorganized and invasive growths in PBS controls that do not have a recognizable basal layer and do not produce a layer of cornified keratin resembling the stratum corneum (Figure 11A) Further, keratinocytes invade into the muscle layer of control skin (arrows, Figure 11A). In contrast, HMW-HA pretreated HaCaT xenografts form polarized spheroids with no histological evidence of invasion into the dermis but rather form polarized structures with a discernable basal layer (Figure 11A, arrowheads) as well as dense outer cornified layers typical of skin architecture (Figure 11A, dotted arrows). Interestingly, immunofluorescence analyses of proliferation detected by human-specific KI67 is similar in both control and HMW-HA treated HaCaT spheroids (Figure 11B) but HMW-HA-treated HaCaT spheroids express higher levels of CDH1 than controls (Figure 11C) consistent with a retention of polarized skin architecture. CDH1 expression is typically expressed by the epidermal keratinocytes where it functions in adherens junctions to maintain differentiation in normal epidermis^55^]. Immunofluorescence assays using human-specific antibodies for TP53 and KRT15 also reveal an increase in apoptotic cells (Figure 11D) and a decrease in TP53+:KRT15+ keratinocytes (Figure 11E) in HMW-HA pre-treated HaCaT keratinocytes compared with PBS treated controls. To assess the effect of HMW-HA on oncogenic signaling in HaCaT xenografts, paraffin-processed tissue sections were stained for GLI1, whose expression is elevated with activation of UVB-induced SHH signaling (Figure 5A). Mouse skin was exposed to UVB in both control and experimental groups prior to implantation of HaCaT spheroids but HAPE or vehicle alone were not applied. Results show that GLI1 expression is similarly elevated in UVB-exposed normal skin surrounding control and HMW-HA-pretreated HaCaT spheroids (Figure 12A, upper panel). However, HMW-HA pretreatment reduced GLI1 expression compared to PBS controls (Figure 12A, lower panel), like the effects of HAPE in *Hr*^-/-^:*Ptch1*^+/-^ mice and HaCaT keratinocytes in vitro (Figure 5A, B).

**Figure 11.**
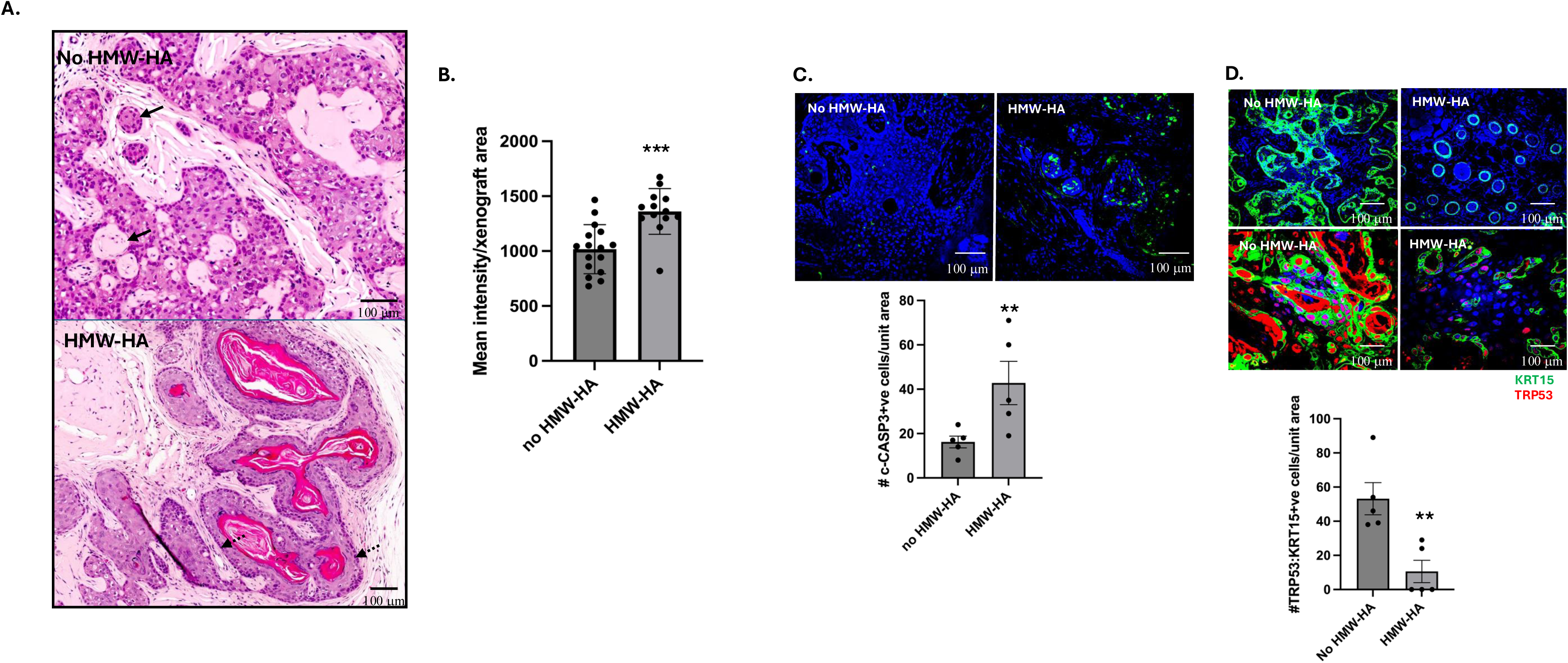
HMW-HA pretreatment increases apoptosis of TP53+:KRT15+ HaCaT subsets *in vivo*. A-C. HaCaT keratinocytes pre-treated with either HMW-HA or PBS (no HMW-HA) were injected intradermally into Nude mice that had been exposed to UVB to generate a mutated cancerized field. **A.** H&E of tissues sections of HaCaT xenografts shows that pretreatment with HMW-HA promotes organization of cells into polarized spheroids that are not invasive, exhibit an organized basal layer of cells (dotted arrows) and produce a cornified layer (red stain). In contrast, PBS-treated controls exhibit disorganized growth patterns, evidence of invasion into the dermis (arrows) and lack an obvious basal cell layer. **B.** HMW-HA treated HaCaT xenografts express increased expression of the polarity marker, CDH1, relative to PBS controls. **C.** HMW-HA pretreatment significantly increases the number of apoptotic HaCaT keratinocytes, detected by cleaved CASP3 staining. **D.** HMW-HA reduces the number of TP53+ve HaCaT keratinocytes. Immunofluorescent intensity for cleaved CASP3, KRT15 and TRP53 was quantified by Image J. Values are the Mean and SEM. *p<0.05, **p<0.01.

**Figure 12.**
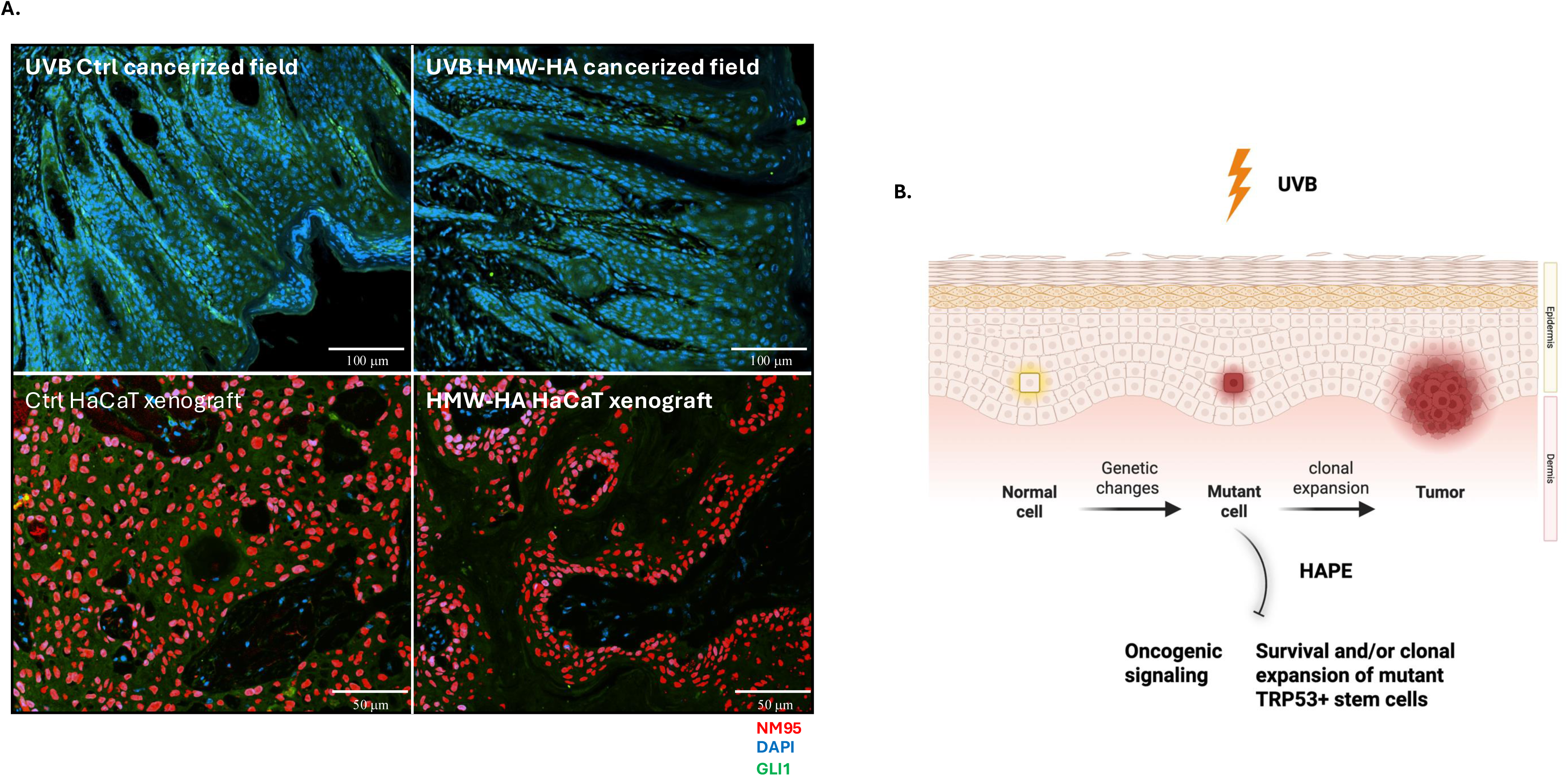
HMW-HA suppresses GLI1 expression in HaCaT xenografts. A. Upper panel: GLI1 immunofluorescence shows protein expression is similar in UVB-exposed normal skin of mice that bear control (Ctrl) or HMW-HA pretreated HaCaT xenografts. These results indicate a similar level of field cancerization surrounding xenografts as indicated by activation of SHH signaling, which results in elevated GLI1 protein expression. Lower panel: Ctrl HaCaT xenografts express higher levels of GLI1 protein expression than HMW-HA-pretreated comparators. B. Model summarizing results of the present study. HAPE promotes the loss of KRT15+ stem cell subpopulations that bear mutant TP53 and suppresses expression of critical oncogenic driver genes, which are required for expansion of mutant cells into tumors.

## DISCUSSION

The collective results of this study are summarized in the Model (Figure 12B) which shows that although oncogenic mutations are established, HMW-HA suppresses both oncogenic signaling and survival or clonal expansion of KRT15+ stem cells that bear a mutant *TP53* and appear to be required for tumor initiation (Figure 12B). Our results confirm previous evidence that HMW-HA is a tumor suppressor^56^] and further identify novel mechanisms for this anti-tumor function. We used a multi-omics approach to probe the anti-tumor mechanisms of HMW-HA and its topical formulation. We show that this polysaccharide supports retention of normal skin polarity following UVB insult, acting downstream of UVB-induced mutations to both block expression of key oncogenic driver pathways and promote loss of KRT15+ cancer-initiating subsets that bear a mutant *Trp53* (Figure 12). These results provide support for the role of cellular polarity and tissue architecture as tumor suppressors^57–59^] and evidence that extracellular microenvironments and cellular contexts can be barriers to the transforming effects of oncogenic mutations. A major effort in modern oncology seeks to design treatments based on the genes that are somatically mutated in cancers. The evidence for oncogenic mutations in normal tissues^17^] supports this focus, which our results extend to include specific mutations in rare stem cell subsets, which are generally considered to be candidate tumor initiators with their rarity providing a barrier to tumor initiation^20,60, 61^]. Our conclusions are consistent with the critical importance of factors specifically hyaluronan and KRT15+ stem cells that act downstream of mutations as barriers to tumor initiation. Our results also begin to fill a gap in current knowledge of the mechanisms required for somatic mutations to initiate carcinogenesis. Ultimately, results reported here have potential utility in identifying novel therapeutic approaches for preventing tumor initiation and for treating established cancers using add back paradigms regardless of the nature of accumulated somatic mutations.

Somatic mutations including oncogenic driver mutations are relatively common in normal tissues^62^]. In many organs, these driver mutations overlap between normal and malignant tissues, suggesting that the earliest events in cancer evolution are already detectable in normal tissues. Due primarily to its constant exposure to UV and age, normal skin harbors the highest somatic mutation burden, which are carried by keratinocytes, melanocytes and fibroblasts^62].^ The predominant mutation signature is UVB and most of the somatic mutations in normal skin are functionally inconsequential. However critical oncogenic mutations that are consistently reported are *NOTCH*, *FAT* and *TP53*^62^]. Mutant clones with these and other driver mutations can expand under positive selection pressure and either promote or restrict carcinogenesis^63^]. Our results are consistent with these previous skin analyses by showing a typical mutation profile in known oncogenes and a predominant UVB mutation signature for cSCC and cBCC in the UVB-exposed skin of control and HAPE-treated animals. The lack of detectable differences in these genomic properties between control and HAPE-treated skin is consistent with an action of HMW-HA downstream of mutations, as suggested by the reduced expression of oncogenic pathway members via CD44 signaling. Our results also predict that a key downstream factor in blocking tumor initiation is also HAPE-mediated loss of KRT15+ cancer-initiating subsets bearing mutant TRP53. Our co-localization results indicate that at least part of this loss is due to KRT15+:TRP53+ stem cell apoptosis. However, we cannot rule out an additional prominent role for HAPE in restricting clonal expansion of subsets bearing multiple oncogenic mutations as a contributor to its anti-cancer function. Although spatial transcriptomics has the advantage of providing a snapshot of cell subset expression profiles in their intact microenvironments, the sensitivity of this platform limits detection of small clones. The WES analysis performed in the present study is not deep sequencing and so only detects mutations that are in high abundance in a bulk tissue sample. WES also does not provide data on variant allele frequency or a metric for clonal size and dynamics. Analyses over time using RNAseq and spatial genomics with single cell sequencing are required for detecting and characterizing small mutant clones whose expansion may be altered by HAPE. These analyses will also help to establish if HAPE alters the growth of small clones that bear multiple mutant oncogenes in addition to *Trp53* reported in the present study that are necessary for keratinocyte tumor initiation.

Keratinocyte tumors arise in a large field of mutations induced by chronic UV exposure, which results in the independent initiation of multiple tumors and fuels tumor recurrence^64^]. Recent evidence suggests that development of keratinocyte tumors within such a mutant field requires not only direct UVB-induced mutagenesis but changes in the cutaneous microenvironment that include chronic inflammation, immune dysregulation and alterations in microbiota^64^]. Additionally, factors including UVB^64,65],^ that affect expansion of stem or progenitor cells contribute to maintaining a field of mutations and promoting carcinogenesis within that field. All together, these factors select for and promote the growth of malignant clones that outcompete normal counterparts. Our results show that HAPE has multiple effects that alter the oncogenic potential of a field of mutations including broadly suppressing the activation of oncogenic signaling, reducing inflammation and suppressing survival of known cancer-initiating stem cells within the mutated field.

The nature of cBCC and cSCC cancer cells of origin has been controversial, likely as a result of the complex mosaicism of skin and the diverse experimental models used to detect these cells^43^]. Previous studies predict that keratinocyte cancers either originate from stem, but not progenitor cells^66^], or from the dedifferentiation of IEF keratinocytes under the influence of carcinogens into a stem-like state^67^]. Currently, experimental models support a role for basal cell layers of the IEF and hair follicle stem cells as cancer-initiating^20^]. For example, KRT14 and 15, which are typical stem cell markers used to detect basal and hair follicle stem cells respectively, label cells in tumors formed in response to carcinogens including UV radiation^43^]. Our results show that while HAPE targets both KRT14 and 15+ stem cells, its ability to promote loss of KRT15/mutant TRP53+ stem cells in the hair follicle predicts that these cells are critical for tumor initiation. Co-staining of hair follicle KRT15+ stem cells with apoptosis markers predicts that HAPE initiates death of these critical subsets. However, our results do not exclude the possibility that HAPE limits clonal expansion of subsets required for tumor initiation and furthermore does not differentiate between the possibilities that KRT15+ subsets that are lost with HAPE treatment contain or support cancer cells of origin. For example, a previous study shows that KRT14+ cells near but not derived from hair follicles are responsible for forming invasive keratinocyte tumors^65^]. The HAPE-sensitive hair follicle cells identified in the present study could produce factors that are required for expansion of the stem cells originating from the IFE. Resolution of these possibilities requires single cell RNAseq and FACS-assisted capture of the HAPE-targeted subsets for a direct assessment of their tumorigenic potential.

Our results predict the regulation of TRP53 expression and targeted loss of mutant TRP53+ KRT15+ stem cells are key factors in the tumor suppressing functions of HAPE. Results from experimental models have shown the critical role of TRP53 expression in both cSCC and cBCC. *Trp53* loss-of-function accelerates KRT15+-induced cBCC and cSCC growth in experimental models^21,66^], is required for escape of cBCC from dormancy^37^], and increases growth of IEF stem cells that give rise to UVB-induced cBCC and cSCC^38^]. Previous studies have shown that NMR-HA attenuates TP53 activation or promotes its expression depending upon the cellular context. For example, Trp53 activity is attenuated in lung fibroblasts *in vitro* by NMR-HA, which is cytoprotective against stress-induced cell-cycle arrest and death^40^]. However, NMR-HA promotes TP53 expression and consequent *in vitro* and *in vivo* apoptosis of human triple negative breast cancer cells^6^]. Our results offer an *in vivo* reconciliation of these seemingly opposing previous results by showing that the effect of HMW-HA on TP53/Trp53 expression is spatially contextual and dependent on its mutation status. We show that HAPE reduces TRP53 activity judging from the down regulation of its target genes, many of which are also downregulated by NMR-HA in stressed lung fibroblasts. Although HAPE also reduces TRP53 expression in hair follicles and should therefore be cytoprotective, it promotes apoptosis in a subset of hair follicle KRT15+ stem cells. Both triple negative breast cancer cell lines (4T1, BT549)^6^] bear TP53 mutations^68^](ATCC). Based upon these results and those of our present study, we propose that HAPE (and NMR-HA) protect keratinocytes expressing wildtype TRP53 by attenuating its activity/expression but either promote death or restrict growth of cells bearing mutant TRP53, particularly gain-of-function mutations (Figure 12). For example, UVB exposure results in loss- and gain-of-function TP53 mutations^69,70^], and knockdown of the expression of *TP53* bearing gain-of-function mutations in the HaCaT keratinocyte line promotes cell death^51–53^]. We tested this possibility by exposing these keratinocytes to HAPE and demonstrating that this causes death of a KRT15 subsets but not stem cells marked by KRT14. HaCaT keratinocytes form cSCC growths when implanted as Matrigel spheres in Nude mice^71^], producing disorganized cell clusters that invade into muscle. The ability of HMW-HA *in vitro* pretreatment to promote the death of KRT15/mutant TP53+ subsets and acquisition of a polarized architecture in *in vivo* spheroids with discernable basal and cornified layers that do not invade the muscle layer supports our proposal that these cells are cancer-initiating and are targeted by HMW-HA. TP53 has previously been implicated in contributing to homeostatic tissue by constraining stem cells and acting as a barrier to dedifferentiation and transcriptome reprograming^72^]. Our results show that HMW-HA is a critical regulator of TP53 expression that particularly regulates its functions in tissue architecture to provide cancer resistance.

Cutaneous keratinocyte tumors are clinically important since they represent 1/3 of cancers diagnosed worldwide^73^], their incidence is increasing and they frequently recur. Current treatments include targeted and non-specific approaches, but these have not been particularly effective in reducing recurrence^13,14^]. Although cSCC and cutaneous basal cell carcinoma (cBCC) are generally not lethal, they are a large health care burden that might be reduced by identifying tumor-resistance mechanisms. These keratinocyte tumors are ideal for probing the mechanisms of tumor initiation and clonal expansion in cancerized fields since the dominant carcinogen, UV, creates a large mutation burden in skin, which is well-characterized for its mutation signatures including oncogenic driver genes ^15,16^]. Our results provide insight into the roles of HMW-HA in maintaining skin homeostasis as a barrier to cBCC and cSCC tumor initiation that could guide therapeutic intervention into these and other cancers.

---

**S. Figure 1.**
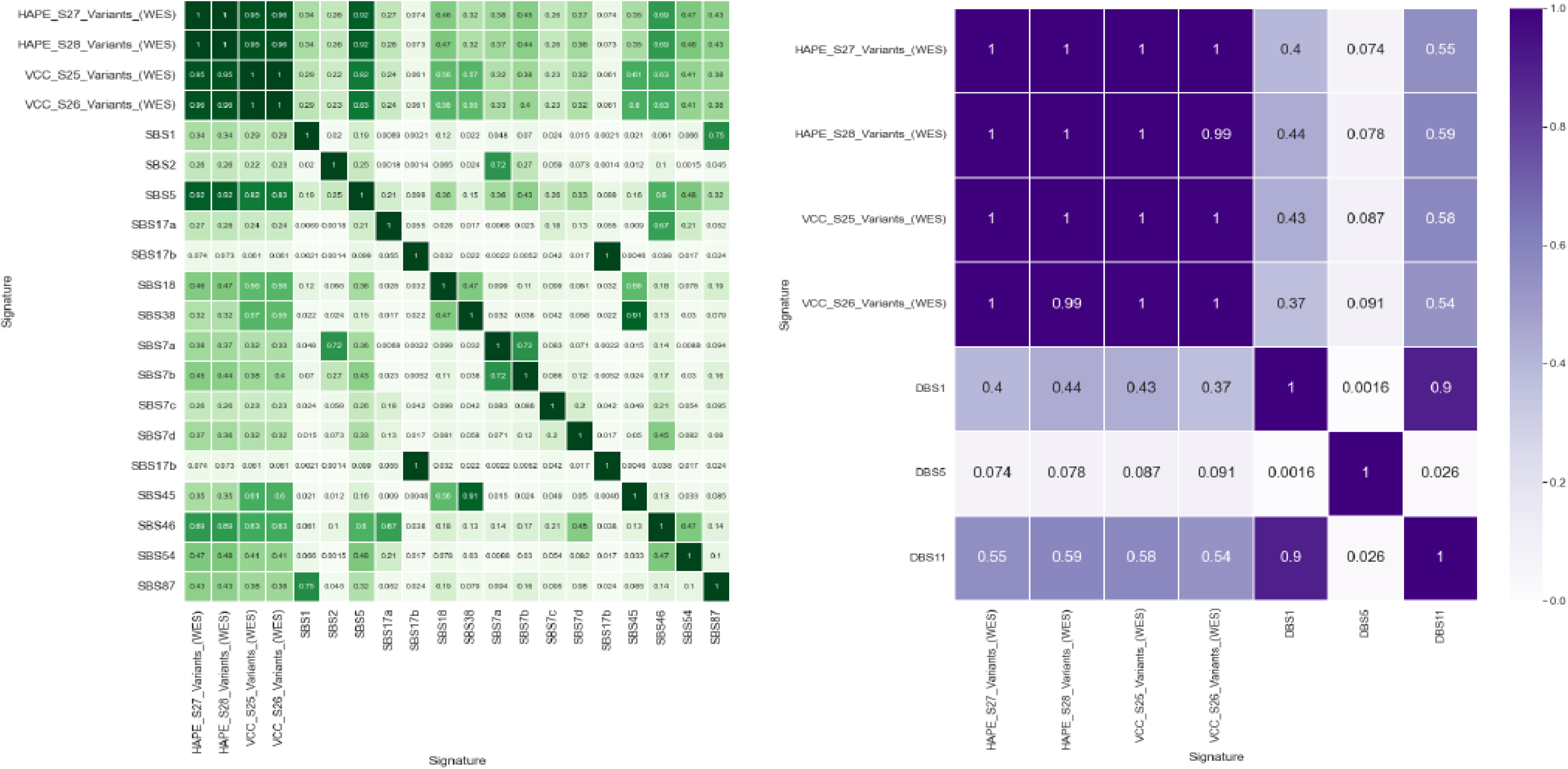
Single base substitution signature plots of HAPE and vehicle mutations are similar. Single base substitution (SBS) mutational signature plots were compared to COSMIC SBS mutational signatures (https://cancer.sanger.ac.uk/cosmic), particularly those generated by UV. Results show a high degree of similarity between the mutational signatures of UVB-exposed HAPE and vehicle control samples.

